# Integrative isotopic Paleoecology (δ^13^C, δ^18^O) of a Late Pleistocene vertebrate community from Sergipe, NE Brazil

**DOI:** 10.1101/482752

**Authors:** Mário André Trindade Dantas, Alexander Cherkinsky, Carlos Micael Bonfim Lessa, Luciano Vilaboim Santos, Mario Alberto Cozzuol, Érica Cavalcante Omena, Jorge Luiz Lopes da Silva, Alcides Nóbrega Sial, Hervé Bocherens

## Abstract

Isotopes are one of the best tools to reconstruct the Paleoecology of extinct taxa, yielding insights about their diet (through carbon; C_3_ and C_4_ plants), niche breadth (*B*_*A*_) and the environment in which they lived. In the present work we go deeper in the use of isotopes and explore a mathematical mixing model with the stable isotopes of two elements (carbon and oxygen) to (1) suggest the relative contribution of four types of food resources (leaves, fruits, roots and C_4_ grass) for meso- and megaherbivores (weight > 100 kg) that lived in the Late Pleistocene of Poço Redondo, Sergipe, Brasil, and (2) evaluate which of these herbivores could be the potential prey for the carnivores *Smilodon populator* and *Caiman latirostris*. To explore the intra/interspecific competition of these fauna, we generate weight estimation, standardized niche breadth (*B*_*A*_) for the meso-megamammals from Sergipe and compare with data from the meso-megaherbivores from Africa, concluding that *Eremotherium laurillardi* and *Toxodon platensis* were the best resource competitors in the Late Pleistocene of Sergipe, and reinforcing their importance as key species in this extinct community. Finally, we reconstructed the paleoenvironment in which the vertebrate community of Sergipe lived, estimating Mean Annual Temperature (°C), Mean Annual Precipitation, Biomass and Energy Expendidure, noting that environments in the Late Pleistocene of Sergipe were similar to those of Africa nowadays, but hotter and with more energy expenditure for these meso-megamammals.

## 1. Introduction

During the last decades, isotopes have been used in Palaeoecology to infer diet of extinct (and extant) taxa, based primarily in carbon isotopic data (*e.g.* Bocherens *et al.*, 1996; MacFadden, 2005; França *et al.*, 2014a), while nitrogen isotopic data has been used as well to infer carnivory or omnivory in mammals (*e.g.* Bocherens *et al.*, 2016). The isotopic approach represented a major advance in paleoecological studies, helping to infer two main food resources for herbivores (C_3_ and C_4_ plants) and paleoenvironmental reconstruction in which herbivores and carnivores could lived (forested or open environments; Kigston & Harrison, 2007; Nelson, 2013; Dantas *et al.*, 2017).

However, isotopes can provide more ecological information than previously thought, such as estimates of niche width and overlap, helping to better understand the ecology of extinct taxa, resource competition and key species in extinct communities (*e.g.* Codron *et al.*, 2007; Dantas *et al.*, 2017), or using two isotope pairs in mathematical mixing models to suggest more than two food resources for herbivores (for example seven resources; Phillips, 2012 and references therein).

Most researchers use carbon and nitrogen isotopic data, extracting these data from collagen. However, in tropical regions these proteins are difficult to be preserved, leaving only the possibility to use carbon and oxygen isotopic data extracted from hydroxyapatite. This mineral usually survives much better than the organic fractions of collagen (Cherkinsky, 2009), being in tropical regions the best option to recover diet information from extinct species.

The isotopic composition of hydroxyapatite can be preserved with minimal or no significant diagenetic alteration. Hydroxyapatite carbonate and phosphate in bone and dentin are more susceptible to diagenetic overprinting than enamel (Bocherens *et al.*, 1996), for example.

Substitutions are mainly in the phosphate position and are most likely in the hydroxyl position. The absorbed carbonates are more labile, but substitute ones are structural carbonates, and, thus, contribute to saving the original isotopic composition (Cherkinsky, 2009).

Thus, the main aims of this paper were to use mathematical mixing models using carbon and oxygen isotopic (extracted from hydroxyapatite) data from fossil vertebrates: (i) to infer four types of resources for herbivores (leaf, fruit, root and C_4_ grass); (ii) to suggest, among the herbivorous mammals from Sergipe, Brazil, which contributed to the isotopic diet of predators such as *Smilodon populator* and *Caiman latirostris*; (iii) to suggest a trophic web structure for the Sergipe community during the Late Pleistocene; (iv) to infer whom were the better competitors (key species) for food resources among herbivores; and, finally, (v) to suggest a paleoenviromental reconstruction in which these taxa could have lived through the Late Pleistocene of Sergipe, estimating Mean Annual Temperature, Mean Annual Precipitation, Biomass and Energy Expendidure.

## 2. Materials and methods

### 2.1. Dataset

Sixteen samples (Table S1) of adult individuals of *Eremotherium laurilardi* (Lund, 1842) (one exception is LPUFS 5693, assigned to a juvenile; Figure S1), *Catonyx cuvieri* (Lund, 1839), *Pachyarmatherium brasiliense* Porpino, Bergqvist & Fernicola, 2009, *Holmesina paulacoutoi* (Guerra & Marecha, 1984), *Glyptotherium* sp., *Panochthus* sp., *Toxodon platensis* Owen, 1837, *Palaeolama major* (Liais, 1872), *Equus* (*Amerhippus*) *neogeus* Lund, 1840 and *Smilodon populator* Lund 1842 from two localities in Sergipe (Fazenda Charco and Fazenda São José, Poço Redondo; Figure 1) were analyzed to obtain carbon and oxygen isotopic composition from the structural carbonate of their bones, dentin and enamel. The samples were collected in “tanks”, which are natural depressions on Neo-Mesoproterozoic lithotypes, characterized by numerous fractures as a result of physical and chemical erosion, and contain sediments transported by seasonal rains, including the remains of animals and plants accumulated during the dry season. The sediments in these depressions are estimated to be of Late Pleistocene and Holocene ages.

**Figure 1.**
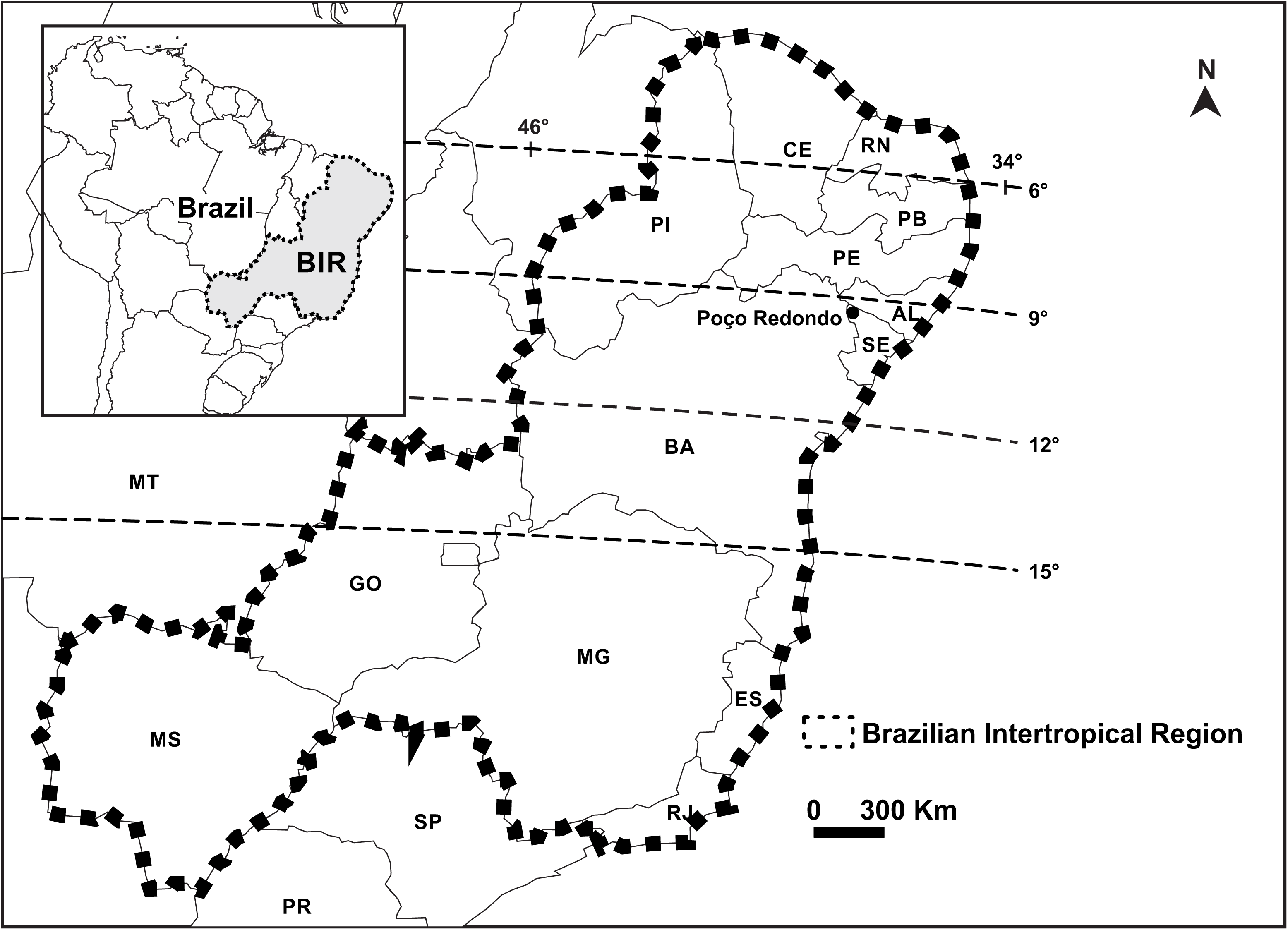
Brazilian Intertropical Region map (*sensu* Oliveira *et al.*, 2017) showing Poço Redondo municipality in Sergipe state, Brazil.

The stable isotope analyzes were performed at “Departamento de Geologia” in “Centro de Tecnologia e Geociências” of Universidade Federal de Pernambuco (Recife, Pernambuco, Brazil) and in Center for Applied Isotope Studies from University of Georgia.

All samples were cleaned by ultrasonic bath with distilled water and then left to dry naturally. The samples were then crushed into smaller fragments to be treated with diluted 1N acetic acid to remove surface absorbed and secondary carbonates. Periodic evacuation ensured that evolved carbon dioxide was removed from the interior of the sample fragments, and that fresh acid was allowed to reach even the interior micro-surfaces.

The chemically cleaned samples were then reacted under vacuum with 100 % phosphoric acid to dissolve the bone/dentine/enamel mineral and release carbon dioxide from hydroxyapatite. The resulting carbon dioxide was cryogenically purified from other reaction products and catalytically converted to graphite (Cherkinsky, 2009).

Graphite ^14^C/^13^C ratios were measured using a CAIS 0.5 MeV accelerator mass spectrometer. The sample ratios were compared to the ratio measured from the Oxalic Acid I (NBS SRM 4990). The ^13^C/^12^C ratios were measured separately using a stable isotope ratio mass spectrometer with respect to PDB.

All results are reported using delta notation, δ = [(R_sample_/R_standard_ - 1)*1000] (Coplen, 1994). The reference for carbon isotope values (R = ^13^C/^12^C) is V-PDB, and oxygen isotope values (R = ^18^O/^16^O) is V-SMOW.

The studied samples were not dated, however we noticed that in Fazenda São José, Poço Redondo, Sergipe has many ^14^C AMS and Electron Spin Resonance - ESR - datings for *Eremotherium laurillardi* and *Notiomastodon platensis* (Table S1) which allow us to suggest that this fossil accumulation had relatively low amount of time-averaging (~32 ky, considering both ^14^C AMS and Electron Spin Resonance datings techniques), similarly to another fossil assemblage in northeastern Brazil (~59 ky, based only in ESR datings; Baixa Grande, Bahia; *e.g.* Ribeiro *et al.* 2014). In addition, França *et al.* (2014a) reported that isotopic diet (δ^13^C) of *Eremotherium laurillardi* and *Notiomastodon platensis* did not changed between 12-19 ky, suggesting a stable environment.

### 2.2. Additional published data

In order to complement our results, and refine the determination of the isotopic diet of Sergipe taxa, we included previously published isotopic data (δ^13^C and δ^18^O, of which most have ^14^C AMS datings; Table S1) and ^14^C AMS and ESR datings of *E. laurillardi*, *N. platensis*, *T. platensis* and *Caiman latirostris* (Dantas *et al.*, 2011; França *et al.*, 2014b; Dantas *et al.*, 2017, and references therein).

Furthermore, we also compared our results and refined isotopic diet data of following extant African mesoherbivores (weight between 100 kg and 750 kg) and megaherbivores (weight > 800 kg) from Kenya and Tanzania (Bocherens *et al.*, 1996; Kingston & Harrison, 2007; Cerling *et al*., 2008): *Loxodonta africana* (Blumenbach, 1797), *Equus quagga* Boddaert, 1785 (= *E. burchelli*), *Diceros bicornis* (Linnaeus, 1758), *Ceratotherium simum* (Burchell, 1817), *Connochaetes taurinus* (Burchell, 1823), *Syncerus caffer* Sparrman, 1779, *Kobus ellipsiprymnus* (Ogilby, 1833), *Oryx beisa* Rüppell*, 1835, Giraffa camelopardalis* (Linnaeus, 1758) and *Hippopotamus amphibius* Linnaeus, 1758 (Table S3).

### 2.3. Multivariate analyses

Carbon (δ^13^C) and oxygen (δ^18^O) isotopic data for meso-megaherbivores from Poço Redondo (Sergipe, Brasil) and Africa was submited to cluster analyses (Q-mode) using the weight pair group method with simple arithmetic averages (UPGMA), using Euclidean Similarity coefficient. A Bootstrap test (N = 10000) was applied to evaluate the consistency of the clusterings. A Principal Component Analyses (PCA) was made as well. All analyses were performed in PAST 2.17 (Hammer *et al.*, 2001).

### 2.4. Weight estimation

To enhance discussion, we calculated the estimated weight (1) for megafauna species that lived in Sergipe (Table 1, Table S1) using the following regression (Anderson *et al.*, 1985):

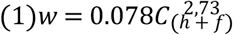

**Table 1.**
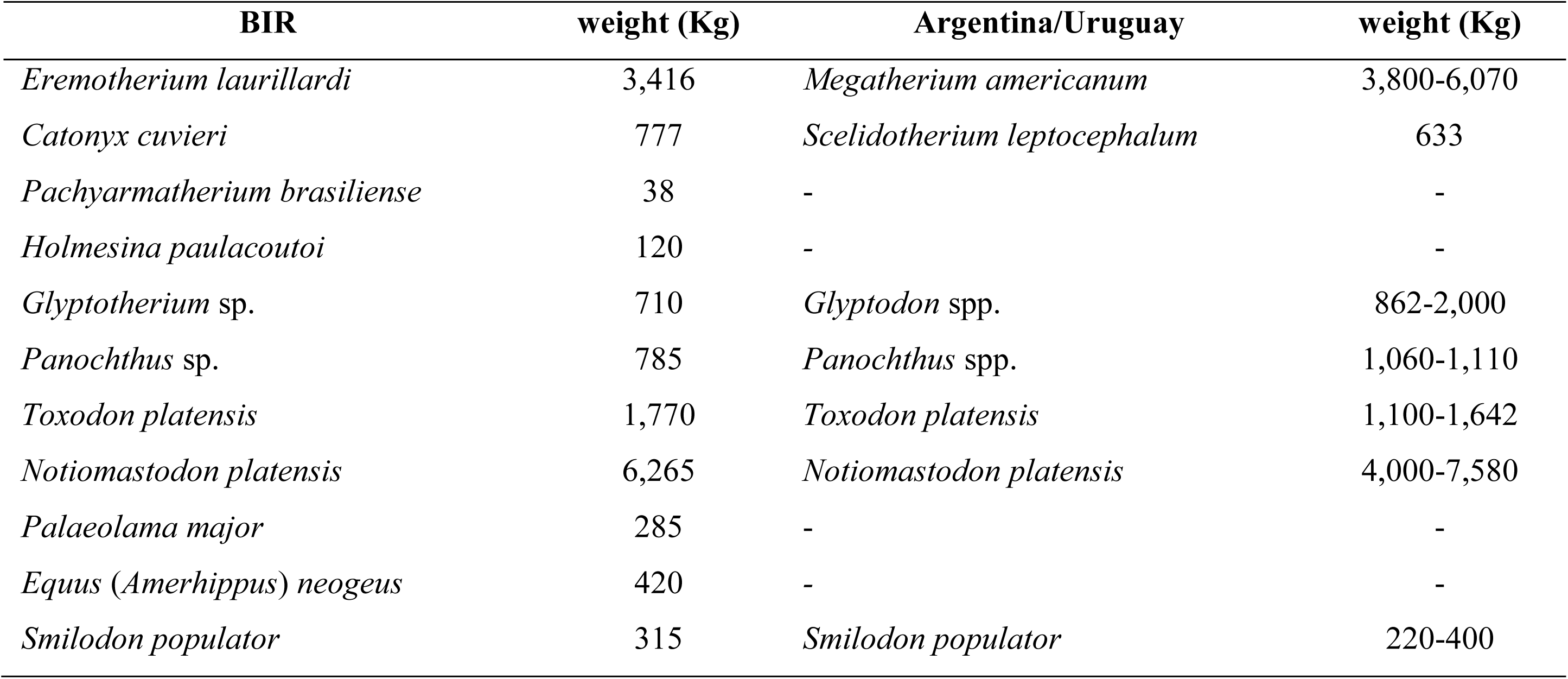
Weight estimation from pleistocenic mammals from Brazilian Intertropical Region, and ir relatives in Argentina and Uruguay (Fariña *et al.*, 1998; Christiansen & Harris, 2005; Prevosti & Vizcaino, 2006

| BIR | weight (Kg) | Argentina/Uruguay | weight (Kg) |
| --- | --- | --- | --- |
| *Eremotherium laurillardi* | 3,416 | *Megatherium americanum* | 3,800-6,070 |
| *Catonyx cuvieri* | 777 | *Scelidotherium leptcephalum* | 633 |
| *Pachyarmatherium brasiliense* | 38 | - | - |
| *Holmesina paulacoutoi* | 120 | - | - |
| *Glyptotherium* sp. | 710 | *Glyptodon* spp. | 862-2,000 |
| *Panochthus* sp. | 785 | *Panochthus* spp. | 1,060-1,110 |
| *Toxodon platensis* | 1,770 | *Toxodon platensis* | 1,100-1,642 |
| *Notiomastodon platensis* | 6,265 | *Notiomastodon platensis* | 4,000-7,580 |
| *Palaeolama major* | 285 | - | - |
| *Equus (Amerhippus) neogeus* | 420 | - | - |
| *Smilodon populator* | 315 | *Smilodon populator* | 220-400 |

Where *W* is the weight (g), *C* is the minimum circumference of humerus and femur diaphysis (in mm). We calculated an average value of circumferences based on the information available in articles and thesis (Cartelle & Abuhid, 1989; Porpino & Bergqvist, 2002; Castro & Langer, 2008; Porpino *et al.*, 2009; Molena, 2012; Oliveira *et al.*, 2017) and in some collections that were accessible (Laboratório de Paleontologia, Universidade Federal de Sergipe; Museu de Ciências Naturais, Pontifícia Universidade Católica de Minas Gerais). When the circumference information was not available, we estimated it using the minimum width of humerus and femur diaphysis as a diameter (*d*) measure (Table S2), through a circumference estimation: *C* = *d*π.

Xenarthrans have flat femur with a high circumference of diaphysis values, leading to an overestimation of weight if using standard method. To avoid this problem, we multiplied their femur circumference by 0.4, trying to acquire a more realistic weight estimation (2). The regression adaptation was calibrated using values for three extant taxa, one Cingulata (*Priodontes maximus* (Kerr, 1792)) and two Tardigrada (*Tamandua tetradactyla* (Linnaeus, 1758) and *Myrmecophaga tridactyla* Linnaeus, 1758) (Table S2). Exceptions were made for gliptodonts (*Panochthus* and *Glyptotherium*) which weight were estimated by original regression proposed by Anderson *et al.* (1985). For extant African megamammals we used maximum estimate weights from Coe *et al.* (1976).

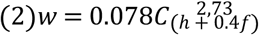

### 2.5. Ecological measurements

To estimate ecological measurements, we calculate isotope niche breadth (*B)* using Levins’ (1968) measure (3), where *p*_*i*_ is the relative proportion of individuals in isotope bin *i*. This measure was then standardized (*B*_*A*_) from 0 to 1 following equation (4), where *n* is total number of isotope bins available. Values lower or equal to 0.5 suggests a specialist, and above 0.5, a generalist.

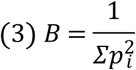

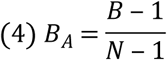

Hence, we also calculated average niche overlaps (*O*) through Pianka’s (1973) index (5), where *p*_*i*_ is relative proportion of individuals in bin *i*. Results between 0 to 0.3 represents low niche overlap; between 0.3 and 0.7, a moderate overlap; and above 0.7, high overlap.

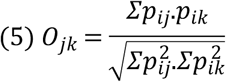

To improve our discussion about niche overlap of Pleistocene megamammals from Sergipe, we estimated population density of herbivores (*D*_*h*_) and carnivores (*D*_*c*_) through Damuth (1981; 1993) general equations (6-7), where we included weight (*w*) in grams:

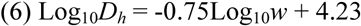

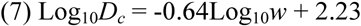

Then we used this information in a modification of Andrades (2018) measurements of intraspecific (*IC*; 8) and interspecific competition (species competition-*SC*; 9) in extinct and extant mammals. Intraspecific competition (*IC*) was calculated dividing population density (*D*) through its standardized niche breadth (*B*_*A*_). Interspecific competition (*SC*) were estimated based on the amount of overlap in isotopic niche between two species in relation to the density of focal species. Values lower or equal to the median were interpreted as low competition, and above, as high competition.

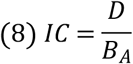

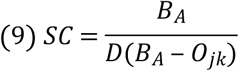

To improve the paleoenvironmental reconstruction, we estimated Energy Expenditure (*Ee*; 10) and Biomass (*Bio*; 11). Through Biomass were estimated secondary production (*Sp*) using ratio of 0.05 to megaherbivores, 0.2 to mesoherbivores and 0.35 to small herbivores from Africa and Sergipe (Coe *et al.*, 1976):

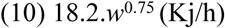

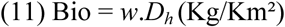

With these informations we could propose Annual Precipitation (*AP*) through regressions (12-14) proposed by Coe *et al.* (1976):

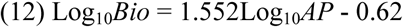

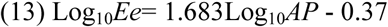

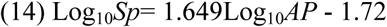

Finally, to estimate the mean and maximum potential prey size for *Smilodon populator*, we used regressions (15-16) proposed for Radloff & Du Toit (2004) for carnivores from Africa:

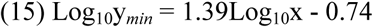

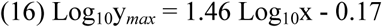

### 2.6. Isotopic diet interpretation using δ^13^C and δ^18^O values

The interpretation of carbon isotopic values for medium-to large-bodied herbivorous mammals are generally made based on known average for C_3_ plants (*μ*δ^13^C= −27±3 ‰), C_4_ plants (*μ*δ^13^C = −13±2 ‰) and CAM plants (intermediate values between δ^13^C of C_3_ and C_4_ plants).

Tejada-Lara et al. (2018) suggested that body mass influences physiological carbon enrichment (□*_diet-bioapatite_) in mammals, and provided equations to determine these values of enrichment. Carbon isotopic data presented here are from mammals (extinct and extant) with body mass varying from 38 to 6,300 kg (Table 1), □*_diet-bioapatite_ varied between 12.47 to 14.84 ‰, we used four values: +12 ‰ for taxa weighting less than 75 kg; +13 ‰ for taxa weighting between 75 kg to 600 kg; +14 ‰ for taxa weighting between 600 kg to 3,500 kg; and, finally, +15 ‰ for taxa weighting more than 3,500 kg (Table S3).

Considering an enrichment of 12-15 ‰, δ^13^C values lower than −15 ‰ to −12 ‰ are typical of animals with a diet consisting exclusively of C_3_ plants, while δ^13^C values higher than −1 ‰ to +2 ‰ are consistent with a diet based on C_4_ plants.

However, C_3_ plants show different values of enrichment. For example, leaves in C_3_ plants are depleted in ^13^C about −1.0 ‰ than others non-photosynthetic tissues like fruits (in average 1.5 ‰) and roots (in average 1.1 ‰), in contrast, C_4_ plants tend to show no enrichment of ^13^C in tissues (*e.g.* fruits, roots) compared to leaves (*e.g.* Yoneyama & Ohtani, 1983; MacFadden, 2005; Cernuzak *et al.*, 2009). Thus we can estimate different type of food resources using carbon isotopic values in association with other isotope in a mathematical mixing model.

In general, nitrogen is used to estimate food resources in a mathematical mixing model with two isotopes (*e.g.* Phillips, 2012, and examples therein), however, analyses in hydroxyapatite are unable to generate nitrogen isotopic data, thus, the only option available is to use oxygen isotopic data instead.

δ^18^O values could be used to help in reconstruction of abiotic conditions, suggesting dry environment (higher ^18^O values) or wet environment (lower ^18^O values; *e.g.* Bocherens & Drucker, 2013), as well, helping to indicate in which guild a vertebrate belongs, because based in its diet, grazers have higher ^18^O values than browsers (*e.g.* Bocherens *et al.*, 1996; Kigston & Harrison, 2007; Cerling *et al*., 2008).

In literature there are no δ^18^O values established for each type of tissue in C_3_ plants (*e.g.* leaves, fruits, roots), only that leaves are more enriched in ^18^O than other non-photosynthetic tissues due to photosynthesis (*e.g.* Cernuzak *et al.*, 2009). A clue to know expected values of δ^18^O in meso-megamammals can be observed in taxa from Africa, in which values between 30-33 ‰ are found in C_4_ grazers, while values near 37 ‰ are found in leaf browsers, as *Giraffa* (Bocherens *et al.*, 1996; Kingston & Harrison, 2007; Cerling *et al*., 2008). An enrichment of δ^18^O in other C_3_ plants tissues can be deduced from results found for forest vertebrates by, for example, Nelson (2013), which shows that fruits (~2 ‰) and roots (~4 ‰) are more enriched than leaves.

Thus, in this paper, we suggest a refinement of proportion that medium-to large-bodied herbivorous mammals (extant; in extinct summing +2 ‰ to compare to modern animals due to Suess effect - Keeling, 1979 - in carbon data) could intake using δ^13^C and δ^18^O values in a two isotopes mathematical mixing model (Phillips, 2012), suggesting as food types: leaves, fruits, roots and C_4_ grass.

Trying to distinguish food resources, we suggest carbon and oxygen isotopic values (Table 2) to be applied in equations (17) in Excel (Microsoft Corporation, Redmond, Washington) through Solver supplement (presuming non-negative values):

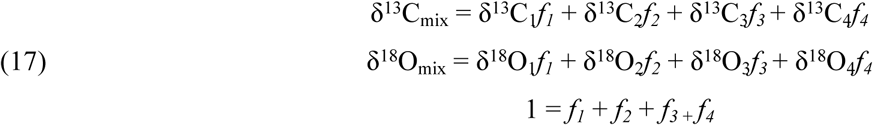

**Table 2.** Carbon (δ^13^C) and oxygen (δ^18^O) isotopic values used in two isotopic mathematical xing model. Carbon values were corrected based on Ɛ^*^_diet-bioapatite_ x weight (*w*) of studied mammals. Oxygen values were corrected due differences found in proboscidean isotopic values in Africa (*Loxodonta africana*) and Sergipe (*Notiomastodon platensis*).

| Food<br>resources | $\delta^{13}\text{C}$ | | | | $\delta^{18}\text{O}$ | |
| --- | --- | --- | --- | --- | --- | --- |
|  | w < 75 Kg | 75 Kg < w < 600 Kg | 600 Kg < w < 3,500 Kg | w > 3,500 Kg | África | Sergipe<br>(+2.5 ‰) |
| **Leaves** | -17.00 | -16.00 | -15.00 | -14.00 | 36.00 | 38.50 |
| **Fruits** | -15.00 | -14.00 | -13.00 | -12.00 | 28.00 | 30.50 |
| **Roots** | -13.00 | -12.00 | -11.00 | -10.00 | 24.00 | 26.50 |
| **C<sub>4</sub> grass** | -1.00 | 0.00 | 1.00 | 2.00 | 32.00 | 34.50 |

For Africa and Sergipe, we used the same isotopic values for carbon (Table 2), however, for oxygen, we used values of Africa as proxy, as this is where the last terrestrial megamammals live there, and apparently in an environment similar to where Pleistocene mammals from South America lived (*e.g.* Cartelle, 1999).

To compare the δ^18^O_CO3_ from Africa with those from Sergipe we considered δ^18^O_CO3_ values of proboscideans as a “thermometer”, since these animals are considered evaporation-insensitive taxa (*e.g.* Yann *et al.*, 2014) and water in plant tissues carries the same ^18^O isotopic signal than source water of environment (Marshall *et al.*, 2007).

Thus, we used the mean values of δ^18^O_CO3_ found in *Loxodonta africana* (Proboscidea; *μ*δ^18^O_CO3_ = 30.03±1.05 ‰; Figure 2A) in comparison with δ^18^O_CO3_ values of *Notiomastodon platensis* (Proboscidea; *μ*δ^18^O_CO3_ = 32.57±1.95 ‰; Figure 2B) as proxy to establish the enrichment (+2.5 ‰) of ^18^O Sergipe environment, and thus, correct δ^18^O_CO3_ values in different tissues of C_3_ plants (leaves, fruits, roots) and C_4_ grass.

**Figure 2.**
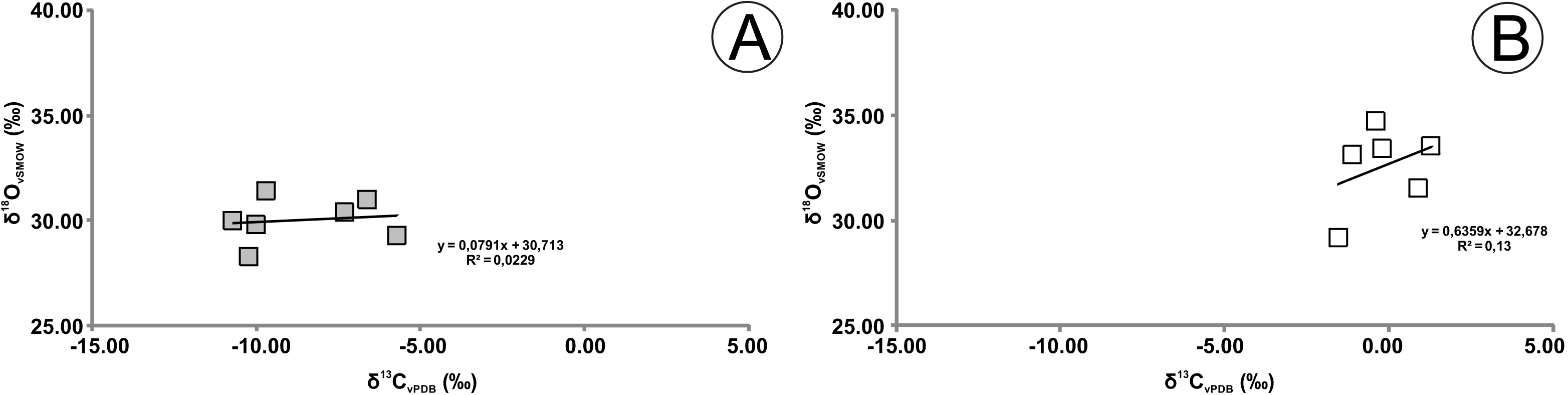
Correlation between δ^13^C and δ^18^O values of (A) *Loxodonta africana* and (B) *Notiomastodon platensis*.

Using δ^18^O_CO3_ of *L. africana* and *N. platensis* we estimate, as well, the Mean Annual Temperature - MAT (°C) in Africa and in the Late Pleistocene of Poço Redondo, Sergipe, Brazil. To do that we considered that δ^18^O_CO3_ is enriched in ~8.7 ‰ than δ^18^O_PO4_ (Bryant *et al*., 1996), then using δ^18^O_PO4_ we estimate δ^18^O_water_ (meteoric water) through regression (18) presented by Ayliffe *et al.* (1992), and, finally, estimated MAT through regression (19) presented by Rozanski *et al.* (1993).

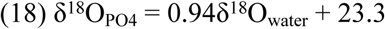

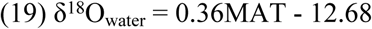

For carnivores (*Smilodon populator* and *Caiman latirostris*) we used the same equations used to estimate food resources for herbivores (11), however including as potential preys the taxa present in Poço Redondo, Sergipe (Tables 3-4), summing +2 ‰ due to Suess effect (Keeling, 1979) and −1 ‰ due to trophic level (Bocherens & Drucker, 2013) in carbon data.

**Table 3.**
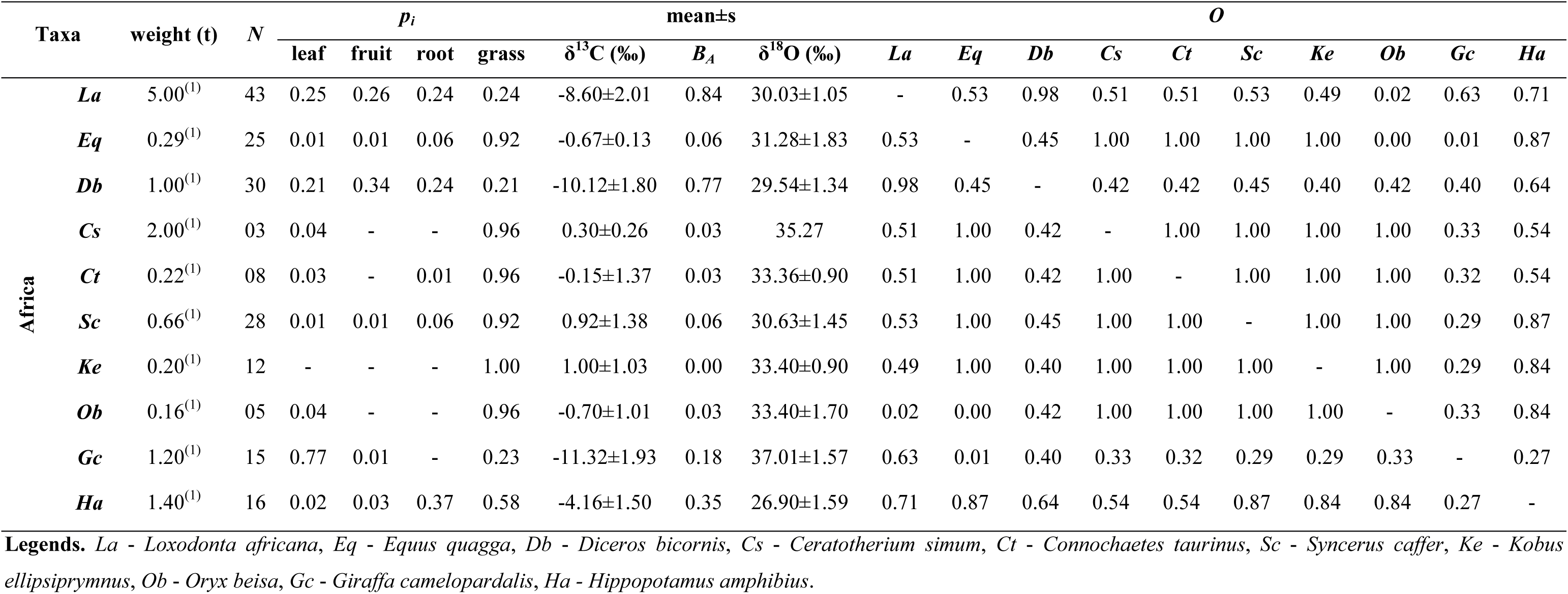
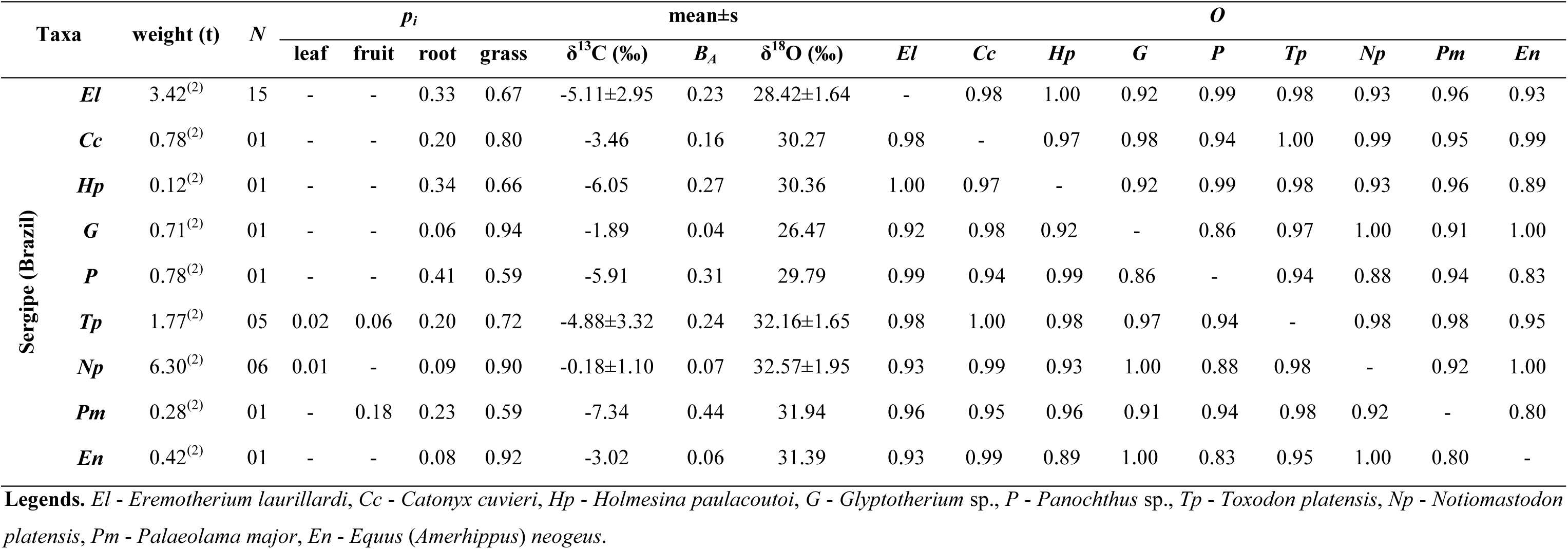
Weight (t), mean values of proportional contributions (*p*_*i*_) of food sources (leaf, fruit, root, C_4_ grass), carbon isotopes (δ^13^C), standardized isotopic niche breadth (*B*_*A*_), oxygen isotopes (δ^18^O), and isotopic niche overlap (*O*) for extant meso-megamammals from Africa and Pleistocene of Sergipe. **References:** ^(1)^ Coe *et al.* (1976); ^(2)^ Our data.

## 3. Results and discussion

### 3.1. Weight estimation

For the Brazilian Intertropical Region (including Sergipe), there are no weight estimations for Pleistocene mammals, thus, following the first attempt proposed by Dantas *et al.* (2017), we continue our effort to know the weight of mammals that lived there, which could help us to better reconstruct the ecology of this fauna.

For localities in Sergipe, we were able to suggest weight measures (Table S2) only for *Eremotherium laurillardi* (*w* = ~3,416 kg) and *Toxodon platensis* (*w* = ~1,770 kg), for further taxa there are no fossils available to estimate their weight. Thus, we estimated weights on the basis of fossils from other geographical regions through Brazilian Intertropical Region (Table S2). Therefore, we estimated and used the following weights: *Catonyx cuvieri* (*w* = ~777 kg), *Pachyarmatherium brasiliense* (*w* = ~38 kg), *Panochthus* sp. (*w* = ~785 kg), *Glyptotherium* sp. (*w* = ~710 kg), *Holmesina paulacoutoi* (*w* = ~120 kg), *Notiomastodon platensis* (*w* = ~6,265 kg), *Palaeolama major* (*w* = ~285 kg), *Equus* (*Amerhippus*) *neogeus* (*w* = ~420 kg) and *Smilodon populator* (*w* = ~315 kg).

In comparison with weight estimated for Pleistocene mammals from Argentina, we note that *E. laurillardi* (*w* = 3,416 kg) is a little smaller than the minimum weight attributed for *Megatherium americanum* (*w* = 3,800-6,070 kg; Table 1). However, this could be related to local environmental conditions (dry environment, see Paleoenvironmental Reconstruction), and not to a flaw in our regression correction. Indeed, we estimated for an *E. laurillardi* in Rio Branco, Acre, using the same approach, a weight of almost 6,600 kg, therefore similar than the maximum weight attributed for *M. americanum*.

The weight of *Catonyx cuvieri* was estimated in 777 kg, which is similar than the weight suggested by Fariña *et al*. (1998) for *Scelidotherium leptocephalum* (median, *w* = 633 kg; Table 1), another South American Scelidotheriinae, which give us confidence in our correction of the regression proposed by Anderson *et al.* (1985).

For Cingulata, we estimated for *Pachyarmatherium brasiliense* a weight of 38 kg. Unfortunately, we did not find any comparable estimation for this taxa. However, it is similar to the weight of the extant armadillo *Priodontes maximus* (*w* = ~19-33 kg), and much smaller than *Holmesina paulacoutoi* (*w* = 120 kg).

The weight of *Glyptotherium* sp. (*w* = 710 kg) was more similar to that suggested for *Glyptodon reticulatus* (*w* = 862 kg) than to *Glyptodon clavipes* (*w* = 2,000 kg; Table 1). Nowadays, specimens of *Glyptotherium* from South America are attributed only to this genus (*e.g.* Oliveira et al., 2010), thus, we estimated the weight for *Glyptotherium texanum* (*w* = 438 kg) and *Glyptotherium arizonae* (*w* = 1,165 kg), based on measures presented by Gilette & Ray (1981), and noticed that *Glyptotherium* from BIR is between proposed weight of this genus.

For *Panochthus* sp., we suggest a weight of 785 kg, which is lower than suggested for Argentinian *Panochthus* spp. (*w* = 1,060-1,110 kg; Table 1); as for *E. laurillardi*, this could be related to local environmental conditions in BIR.

For *Toxodon platensis* (*w* = 1,770 kg), *Notiomastodon platensis* (*w* = 6,265 kg) and *Smilodon populator* (*w* = 315 kg), we suggest similar weights for taxa from Argentina (Table 1); only *Palaeolama major* (*w* = 285 kg) and *Equus* (*Amerhippus*) *neogeus* (*w* = 420 kg) do not have estimations for comparison.

Thus, in Sergipe, we had a fauna composed by one megacarnivore (*S. populator*), one omnivore (*P. brasiliense*), six mesoherbivores (*H. paulacoutoi*, *C. cuvieri*, *Glyptotherium* sp., *Panochthus* sp., *P. major* and *E.* (*A.*) *neogeus*) and three megaherbivores (*E. laurillardi*, *T. platensis* and *N. platensis*).

### 3.2. Isotopic paleoecology (δ^13^C, δ^18^O) of meso-megaherbivores mammals

Attempting to understand better isotopic paleoecology of Sergipe Pleistocene mammals taxa, we reunite here, to compare, available isotopic data for 11 extant mammals from Africa (Table 3 and Figure 3A), to analyze the isotopic diet patterns found through mathematical mixing model using carbon and oxygen. Based in Principal Component Analyses (PCA) we notice that contribution of carbon (*p*_*i*_ = 68 %) is higher than oxygen (*p*_*i*_ = 32 %) in this mathematical mixing model diet refinement, and together with Cluster analyses shows three well defined groups/guilds: browser, mixed-feeders and grazers (Figure 3A-B).

**Figure 3.**
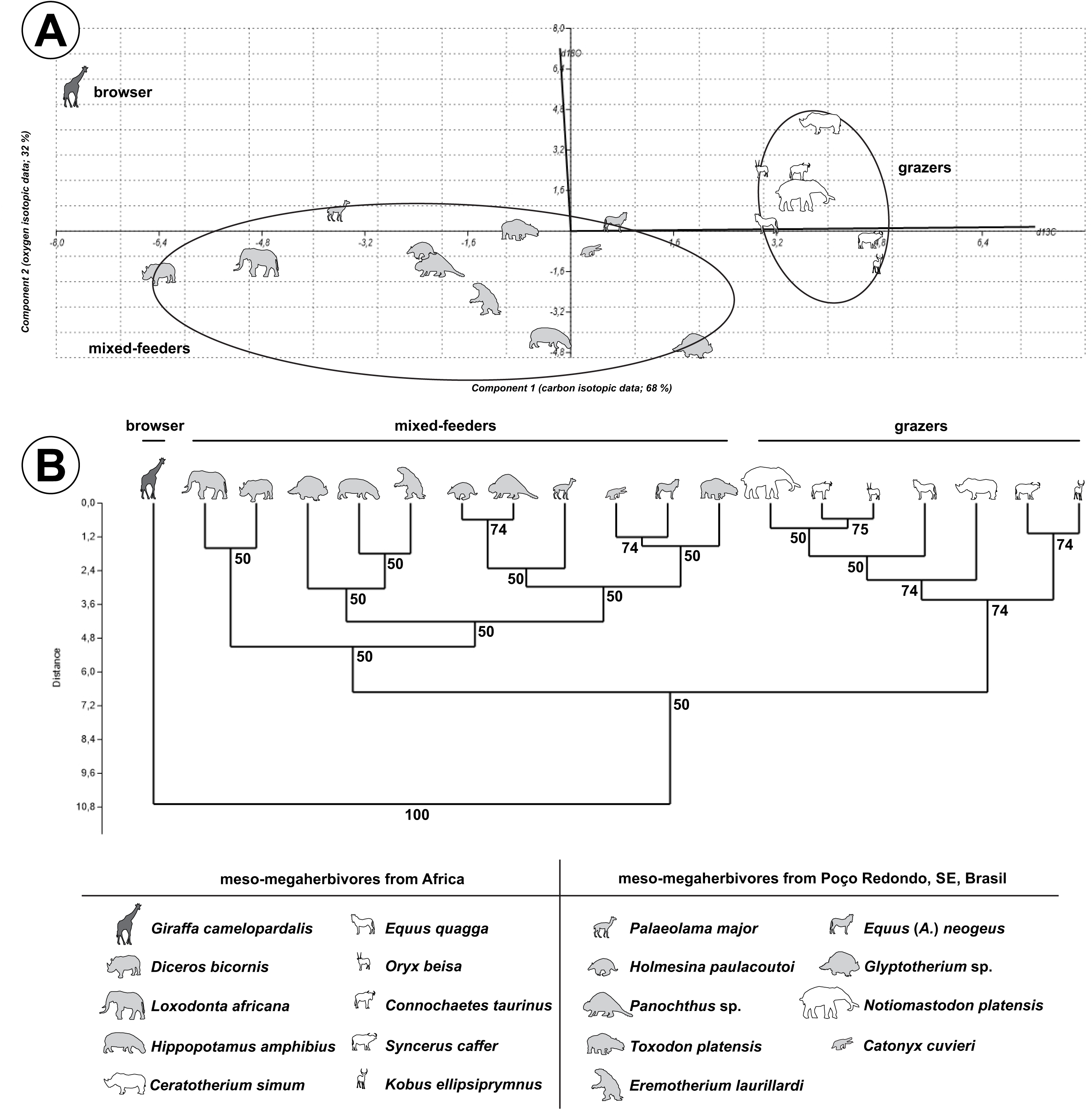
(A) Principal Component Analyses (PCA) using δ^18^O and δ^13^C values for meso-megaherbivores from Africa and Poço Redondo (Sergipe, Brazil); (B) Dendogram with clustering of taxa according to their ecological guilds (Bootstrap test, N = 10.000).

In Africa savannahs there were, at least, 20 species of grazers, 13 of browsers, 10 of mixed-feeders and one of omnivore (Owen-Smith, 1982). Grazer guilds were composed by the megaherbivore *Ceratotherium simum* (*μ*δ^13^C = 0.30±0.26 ‰, *μ*δ^18^O = 35.27 ‰), a specialist grazer (*B*_*A*_ = 0.03) feeding on 96 % of C_4_ grass and 4 % of leaves, and for specialists mesoherbivores *Equus quagga* (*μ*δ^13^C = −0.67±0.13 ‰, *μ*δ^18^O = 31.28±1.83 ‰; *B*_*A*_ = 0.06), *Connochaetes taurinus* (*μ*δ^13^C = −0.15±1.37 ‰, *μ*δ^18^O = 33.36±0.90 ‰; *B*_*A*_ = 0.03), *Syncerus caffer* (*μ*δ^13^C = 0.92±1.38 ‰, *μ*δ^18^O = 30.63±1.45 ‰; *B*_*A*_ = 0.06), *Kobus ellipsiprymnus* (*μ*δ^13^C = 1.00±1.03 ‰, *μ*δ^18^O = 33.40±0.90 ‰; *B*_*A*_ = 0.00) and *Oryx beisa* (*μ*δ^13^C = −0.70±1.01 ‰, *μ*δ^18^O = 33.40±1.70 ‰; *B*_*A*_ = 0.03), which have a diet composed mainly by C_4_ grass (varying from 92 % to 100 %; Table 3; Figure 3A-B), consumption of roots and leaves were low (*p*_*i*_ = 0-6% and 1-4 %, respectively), and fruits (*p*_*i*_ = 0-1 %) virtually inexistent.

In browser guild we have data only for the specialist *Giraffa camelopardalis* (*μ*δ^13^C = −11.32±1.93 ‰, *μ*δ^18^O = 37.01±1.57 ‰; *B*_*A*_ = 0.18) feeding in more than 77% of leaves (Table 3; Figure 3A-B), and, in mixed-feeder guild are *Loxodonta africana* (*μ*δ^13^C = −8.60±2.01 ‰, *μ*δ^18^O = 30.03±1.05 ‰; *B*_*A*_ = 0.84) and *Diceros bicornis* (*μ*δ^13^C = −10.12±1.80 ‰, *μ*δ^18^O = 29.54±1.34 ‰; *B*_*A*_ = 0.77) which fed similarly in C_4_ grass (*p*_*i*_ = 21-24%), roots (*p*_*i*_ = 24%), fruits (*p*_*i*_ = 26-34%) and leaves (*p*_*i*_ = 21-25%; Table 3; Figure 3A-B). *Hippopotamus amphibius* (*μ*δ^13^C = −4.16±1.50 ‰, *μ*δ^18^O = 26.90±1.59 ‰; *B*_*A*_ = 0.35) fed more in C_4_ grass (*p*_*i*_ = 58%), in its case, values attributed to roots (*p*_*i*_ = 37%) could be C_3_ aquatic plants, due to lower isotopic values of carbon and oxygen (Table 3).

For Sergipe, we generated new isotopic data for *Eremotherium laurillardi* and *Toxodon platensis*, plus unpublished isotopic data for *Catonyx cuvieri*, *Holmesina paulacoutoi*, *Glyptotherium* sp., *Panochthus* sp., *Palaeolama major* and *Equus* (*Amerhipus*) *neogeus*, and include published isotopic data for *Eremotherium laurillardi*, *Toxodon platensis*, *Notiomastodon platensis* (França *et al.*, 2014a; Dantas *et al.*, 2017, and references therein; Table 3 and S1; Figure 3A and 4A-B) to refine isotopic diet of all taxa that lived in Sergipe during the Late Pleistocene.

Based in PCA and Cluster Analyses (Figure 3A-B) we notice that only *N. platensis* (*μ*δ^13^C = −0.18±1.10 ‰, *μ*δ^18^O = 32.57±1.95 ‰; *B*_*A*_ = 0.07; *p*_*i*_C_4_ = 90 %) was included in grazer guild, feeding in more than 90% of C_4_ grass, as observed in herbivores mammals from Africa. Remaining studied taxa was included in mixed-feeder guild (Figure 3A-B), being subdivided in four subgroups, which presents mainly variations in consumption of C_4_ plants.

*T. platensis* (δ^13^C = −4.88 ‰, δ^18^O = 32.16 ‰; *B*_*A*_ = 0.24), *E.* (*A.*) *neogeus* (δ^13^C = −3.02 ‰, δ^18^O = 31.39 ‰; *B*_*A*_ = 0.06) and *Catonyx cuvieri* (δ^13^C = −3.46 ‰, δ^18^O = 30.27 ‰; *B*_*A*_ = 0.16) presents major consumption of C_4_ grass (*p*_*i*_ = 72-92 %), but fed well in roots (*p*_*i*_ = 8-20 %). In this subgroup only *T. platensis* presented consumption of leaves (*p*_*i*_ = 2 %) and fruits (*p*_*i*_ = 6 %). The diet of *C. cuvieri* was in contrast to a browser diet expected for a Scelidotherinae (Bargo *et al.*, 2006a, b; Dantas *et al.*, 2017). The analyzed sample (UGAMS 35324; Table S1) belonged to an adult individual (Dantas & Zucon, 2007), thus, we hypothesize that the environmental conditions led to a change of its diet with a major consumption of C_4_ grass and roots.

In other subgroup we have *Palaeolama major* (*μ*δ^13^C = −7.34 ‰, *μ*δ^18^O = 31.94 ‰; *B*_*A*_ = 0.44), *Panochthus* sp. (δ^13^C = −5.91 ‰, δ^18^O = 29.79 ‰; *B*_*A*_ = 0.31) and *Holmesina paulacoutoi* (δ^13^C = 6.05 ‰, δ^18^O = 30.36 ‰; *B*_*A*_ = 0.27), these taxa had a moderate consumption of C_4_ grass (*p*_*i*_ = 59-66 %) and roots (*p*_*i*_ = 23-41 %). In this subgroup only *P. major* presented consumption of fruits (*p*_*i*_ = 18 %). Marcolino *et al.* (2013), through the analysis of coprolites, suggested a diet based on C_3_ plants for this taxon, which is consistent with our results – consumption of 41 % of C_3_ plants tissues (fruits and roots).

Finally, *Eremotherium laurillardi* (*μ*δ^13^C = −5.11±1.95 ‰, *μ*δ^18^O = 28.42±1.64 ‰; *B*_*A*_ = 0.23) and *Glyptotherium* sp. (δ^13^C = −1.89 ‰, δ^18^O = 26.47 ‰; *B*_*A*_ = 0.04) was grouped with *H. amphibius*, mainly because of their lower oxygen isotopic values. For *E. laurillardi* the lower isotopic values of oxygen was attributed to it consumption on roots (*p*_*i*_ = 33 %), while in *H. amphibius* it could be related to its feeding on aquatic C_3_ plants. It is worth noting that LPUFS 5693 was an unworn molariform from a juvenile (perhaps from a suckling individual; Figure S1A-B), and we noticed that the carbon isotopic data (δ^13^C = −6.05 ‰) was not different from the adults of the same locality, being equivalent to mother diet, because carbon isotopic values of mother milk and offspring would not have significant differences (*e.g.* trophic level; Jenkins *et al.*, 2001).

Bargo *et al.* (2006a) suggested that *M. americanum* and *E. laurillardi* had masticatory apparatus with similar biomechanical functions, presenting robust cranial muscles which allow them to have a strong bite, and a thick cone-shaped and prehensile upper lip that could select parts of plants. Following Vizcaino *et al.* (2006) and Bargo *et al.* (2006b) we estimated the occlusal surface area (OSA) and Hypsodonty index (HI) for a dentary of *E. laurillardi* (LPUFS 4755; Figure S1C-D) from Sergipe, to support the discussion of our isotopic results. Estimate OSA from LPUFS 4755 was 10,650.83 mm^2^, which is equivalent for OSA found for *Megatherium americanum* (10,818.36±464.23 mm^2^; Vizcaino *et al.*, 2006), which allow us to suggest that it could process a large amount of turgid to soft food items, as C_4_ grass and C_3_ roots. HI of *E. laurillardi* LPUFS 4755 was higher (HI = 0.96; length of the tooth row = 172.94 mm; depth of the dentary = 166.06 mm) than those found for northeastern Brazilian species (MCL and MNRJ samples; HI = 0.76±0.02; Bargo *et al.*, 2006b), and closer to HI found for *M. americanum* (HI = 1.02±0.07; Bargo *et al.*, 2006b), which is odd, but may suggest in Sergipe a population adapted to a diet with high consumption of dust and grid together with their food items (as C_4_ grass and C_3_ roots), which reinforce our interpretation found while using the mathematical mixing model of carbon and oxygen isotopic values.

For *Glyptotherium* sp. we have only one sample, its lower value of oxygen is odd, as it had a great consumption of C_4_ grass (*p*_*i*_ = 94 %). Gillette & Ray (1981) suggests that *Glyptotherium* could have a semi-aquatic habit, as *H. amphibius*, however, as we have only one sample, we cannot discard or confirm this hypothesis.

### 3.3. Isotopic paleoecology (δ^13^C, δ^18^O) of Sergipe carnivorous vertebrates

In addition to the isotopic data of meso-megaherbivores from Poço Redondo, Sergipe, we generated isotopic data of carbon and oxygen for *Pachyarmatherium brasiliense* and *Smilodon populator*, and included published isotopic data for *Caiman latirostris* (França *et al.*, 2014b), to suggest a trophic web in this assemblage (Figure 5).

Downing and White (1995) suggests for *Pachyarmatherium leiseyi* a diet composed mainly of termites and ants. Unfortunately, as we do not have nitrogen isotopic data for *P. brasiliense*, we could not test this hypothesis. However, following Downing and White (1995), our results in mathematical mixing model (δ^13^C = −6.66 ‰, *μ*δ^18^O = 28.70 ‰; *B*_*A*_ = 0.25) could not reflect a diet on plants, but on insects that collected tissues of these plants, thus, *P. brasiliense* probably fed more on Blattodea (termites and ants) taxa which lived in open areas (69 %; feeding on C_4_ grass).

*Smilodon populator* (δ^13^C = −6.06 ‰, δ^18^O = 30.58 ‰) was a generalist carnivore (*B*_*A*_ = 0.53) and had as main preys (*p*_*i*_ ≥ 10 %; Table 4), *P. major* (14 %), *H. paulacoutoi* (12 %), *Toxodon platensis* (12 %), *P. brasiliense* (11 %), *Panochthus* sp. (12 %), *Catonyx cuvieri* (10 %), *Equus* (*Amerhippus*) *neogeus* (10 %) and *Caiman latirostris* (10 %; Table 4; Figures 4A-5), allowing us to hypothesize that, at least in Sergipe, it did not have a specialization on a prey type, which could suggest a pack-hunting behavior, as individuals of this pack could feed on a variety of preys sampled proportionally.

**Table 4.** Prey isotopic (δ^13^C, δ^18^O) contribution (%) to isotopic diet of two vertebrate carnivores from Sergipe, Brazil.

| Potential preys (weight in Kg) | *Smilodon populator* | *Caiman latirostris* |
| --- | --- | --- |
| *Smilodon populator* (315) | - | - |
| *Caiman latirostris* (60) | 0.10 | - |
| *Eremotherium laurillardi* (3,416) | 0.06 | - |
| *Catonyx cuvieri* (777) | 0.10 | 0.12 |
| *Pachyarmatherium brasiliense* (38) | 0.11 | - |
| *Holmesina paulacoutoi* (120) | 0.12 | - |
| *Glyptotherium* sp. (710) | 0.02 | 0.10 |
| *Panochthus* sp. (785) | 0.12 | - |
| *Toxodon platensis* (1,770) | 0.12 | 0.16 |
| *Notiomastodon platensis* (6,300) | 0.01 | 0.44 |
| *Palaeolama major* (285) | 0.14 | - |
| *Equus (Amerhippus) neogeus* (420) | 0.10 | 0.18 |

**Figure 4.**
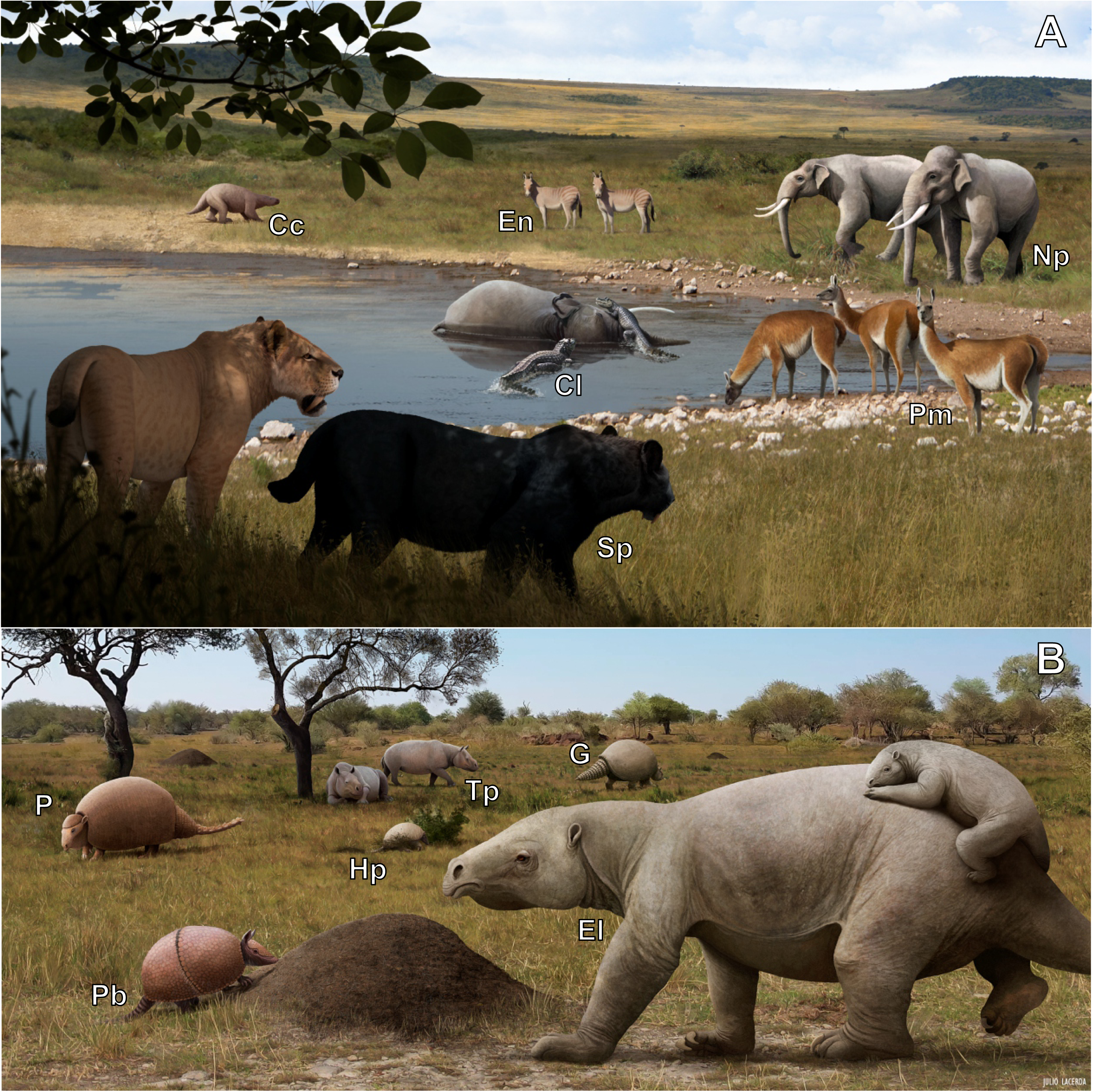
(A) Two *Smilodon populator* stalking a *Palaeolama major* group, while two *Caiman latirostris* are scavenging a *Notiomastodon platensis* corpse; (B) Pleistocene megamammals from Sergipe, Brazil (Image: Julio Lacerda, 2018). **Legends.** Cc - *Catonyx cuvieri*, En - *Equus* (*Amerhippus*) *neogeus*, Np - *Notiomastodon platensis*, Cl - *Caiman latirostris*, Pm - *Palaeolama major*, Sp - *Smilodon populator*, P - *Panochthus* sp., Tp - *Toxodon platensis*, G - *Glyptotherium* sp., Hp - *Holmesina paulacoutoi*, Pb - *Pachyarmatherium brasiliense*, El - *Eremotherium laurillardi*.

**Figure 5.**
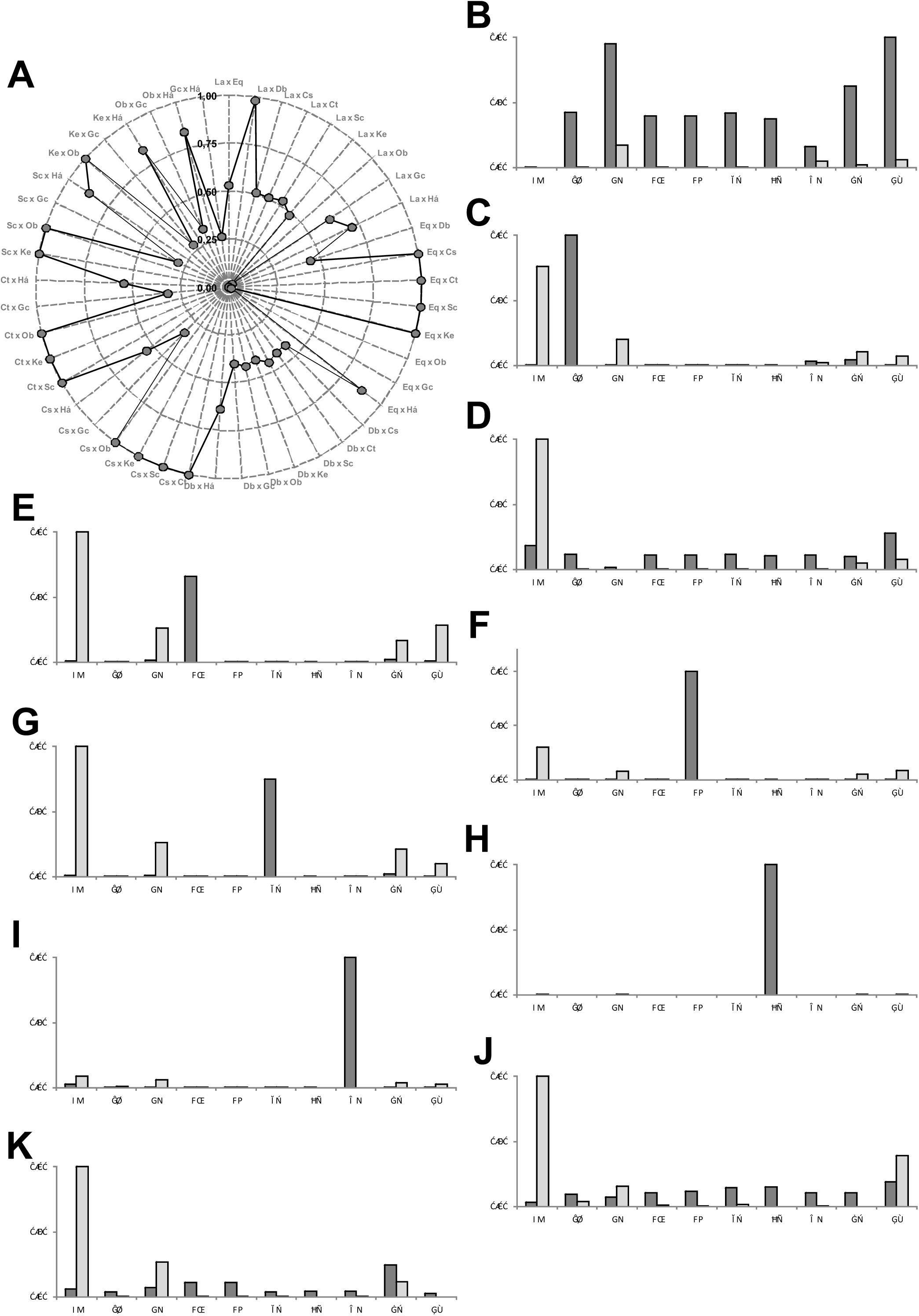
Isotopic (δ^18^O and δ^13^C) trophic web from pleistocenic meso-megamammals from Sergipe, Brazil. **Legends.** bold arrows - food resources that contributed more than 10% in isotopic signature of consumer; thin arrow - food resources that contributed less than 10% in isotopic signature of consumer.

Using regressions proposed by Radloff & Du Toit (2004) we estimate that the mean and maximum prey size for *S. populator* could have varied between 540-3,000 kg, which allows us to suggest that only 36 % of its diet was composed of taxa belonging to his optimal interval (Table 4). Above this limit there is a low percentage (*p*_*i*_ = 7 %), including the megaherbivores *Eremotherium laurillardi* and *Notiomastodon platensis*, however, the majority of its diet (*p*_*i*_ = 56 %) was possibly based on mammals weighting less than 540 kg (Table 4). It is possible that *S. populator* hunted actively *Equus* (*A.*) *neogeus* (*p*_*i*_ = 10 %) and *Palaeolama major* (*p*_*i*_ = 14 %), as their weight is not so distant from their mean prey size. The predation on *C. latirostris* is suggested (*p*_*i*_ = 10 %) based mainly in observation of predation nowadays of the extant Felidae, *Panthera onca* in this taxa (*e.g.* Azevedo & Verdade, 2011). Isotopic contribution of *P. brasiliense* (*p*_*i*_ = 11 %) and *H. paulacoutoi* (*p*_*i*_ = 12 %) for *S. populator* diet could represent a scavenger habits for this carnivore (Table 4).

In southern Chile, Prevosti & Martin (2013), based on carbon and nitrogen isotopes, suggested as possible prey for *Smilodon*: *Hippidion*, indeterminated Camelidae and *Lama guanicoe* (Camelidae), which is similar to our results, as major prey was Camelidae taxa. However, this approach depends on herbivores isotopic data available, and Bocherens *et al.* (2016) suggests as main prey for *Smilodon, Macrauchenia* (Macraucheniidae), followed by *Megatherium* (Megatheriidae) and *Lestodon* (Mylodontidae), which is very different for our results, mainly because we do not have isotopic data for Macraucheniidae and Mylodontinae taxa in Sergipe.

These results could show us a regional difference in prey types for *S. populator*, or, as said previously, be a consequence of the absence of isotopic data for more herbivores in our analyses.

Another carnivore found in Sergipe was *Caiman latirostris* (δ^13^C = −3.01 ‰, δ^18^O = 31.40 ‰; França *et al.*, 2014b), which was more specialist (*B*_*A*_ = 0.27) than *S. populator*, but our analyses allow us to suggest that it could feed on a variety of taxa, mainly on *N. platensis* (44 %), as well, in *E. (A.) neogeus* (18 %), *T. platensis* (16 %), *C. cuvieri* (12 %) and *Glyptotherium* sp. (10 %;Table 4).

We know that *C. latirostris* could not actively hunt mammals taxa found in Sergipe, as they weight more than 420 kg (Table 4), however, we suggest that it could fed on dead corpses, acting as a scavenger like other crocodiles, because this would facilitate dismembering of large prey corpses (*e.g.* Dixon, 1989; Perez-Higareda *et al.*, 1989; Figures 4A-5).

### 3.4. Niche overlap and resources competition in meso-megaherbivores mammals

Africa ecosystems were structured by meso-megaherbivores composed mainly by grazers taxa, belonging to the orders Proboscidea, Cetartiodactyla and Perissodactyla (*e.g.* Cumming, 1982). In our analysis, the grazer guild was composed of one megaherbivore (*C. simum*) and five mesograzers (*E. quagga*, *C. taurinus*, *S. caffer*, *K. ellipsiprymnus* and *O. beisa*), all feeding in more than 90% of C_4_ grass, having a high niche overlap (*O* = 1.00; Table 3). Despite these high values, competition indexes (Table 5) allow us to suggest that intraspecific competition (*IC*; Table 5) was higher than interspecific competition with other grazers (*SC*; Table 5), which was virtual inexistent (Figure 6), probably because they fed on different taxa of C_4_ grass (*e.g.* Arsenault & Owen-Smith, 2011) or acted as resource facilitator for some taxa (*e.g.* Perrin & Brereton-Stiles, 1999), partitioning food resources through body size (*e.g.* Kleynhans *et al.*, 2011).

**Table 5.**
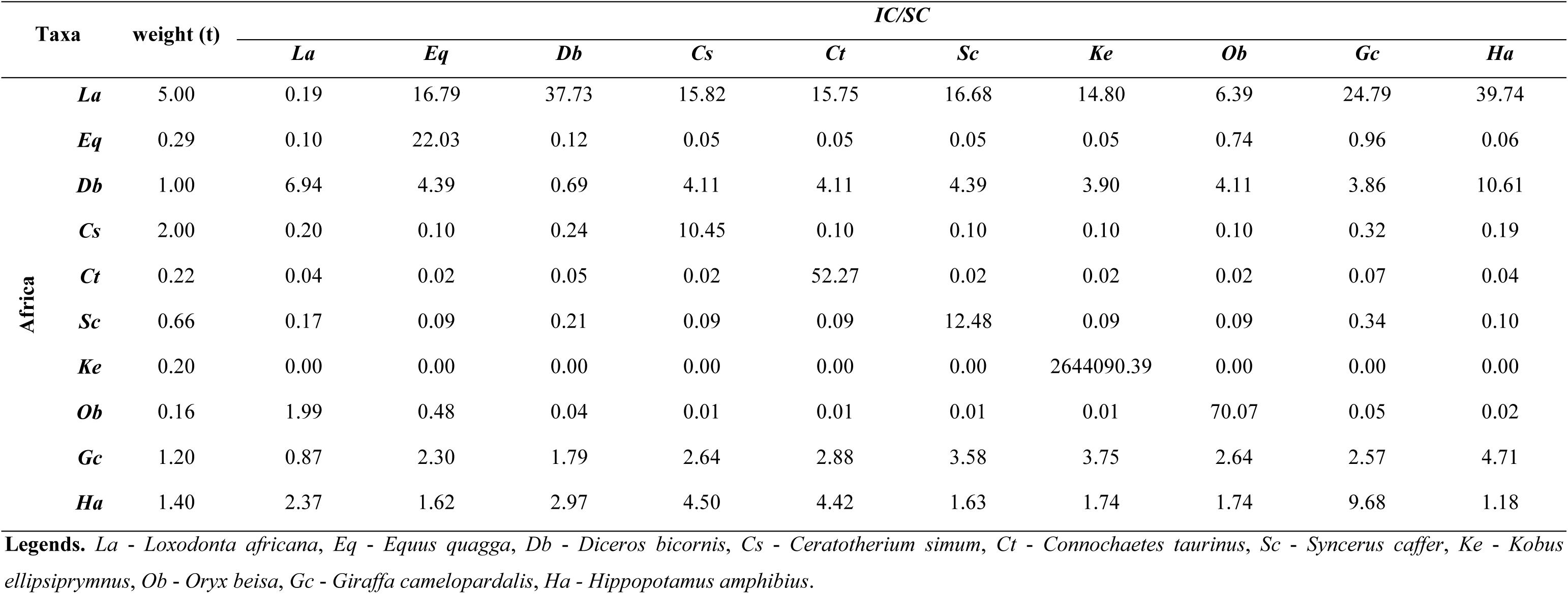
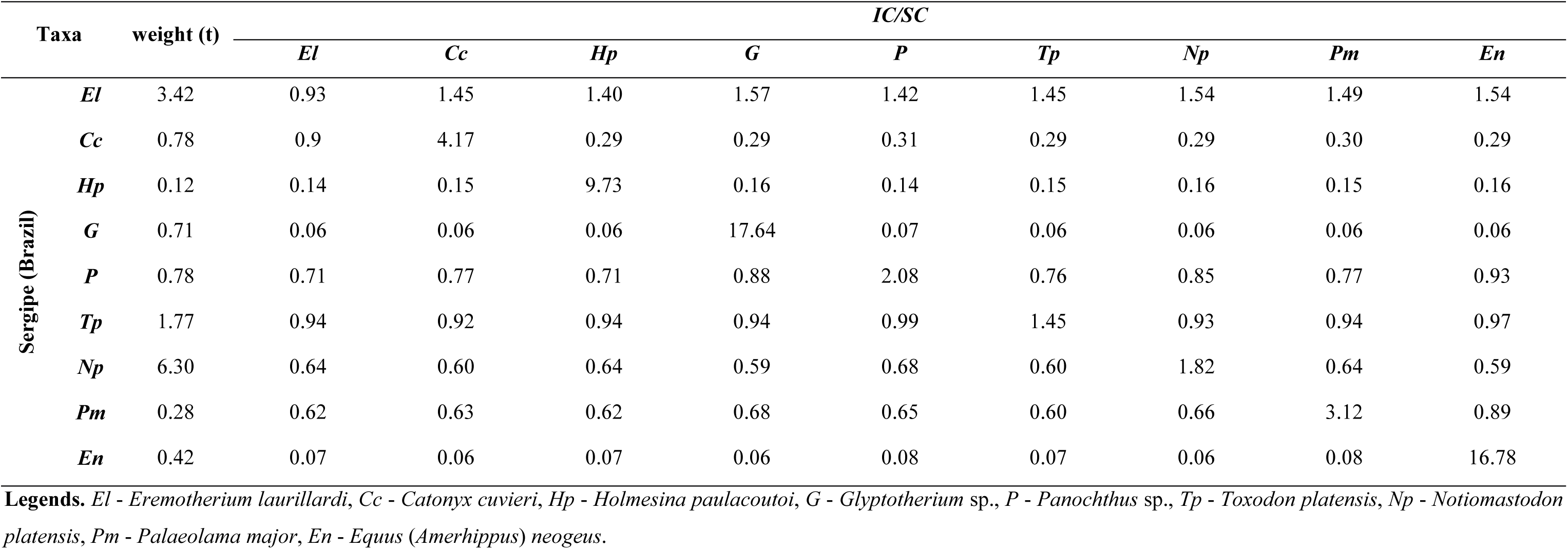
Intraspecific (*IC*) and interspecific (*SC*) competition values for meso-megamammals from Africa and Pleistocene of Sergipe, Brazil.

**Figure 6.**
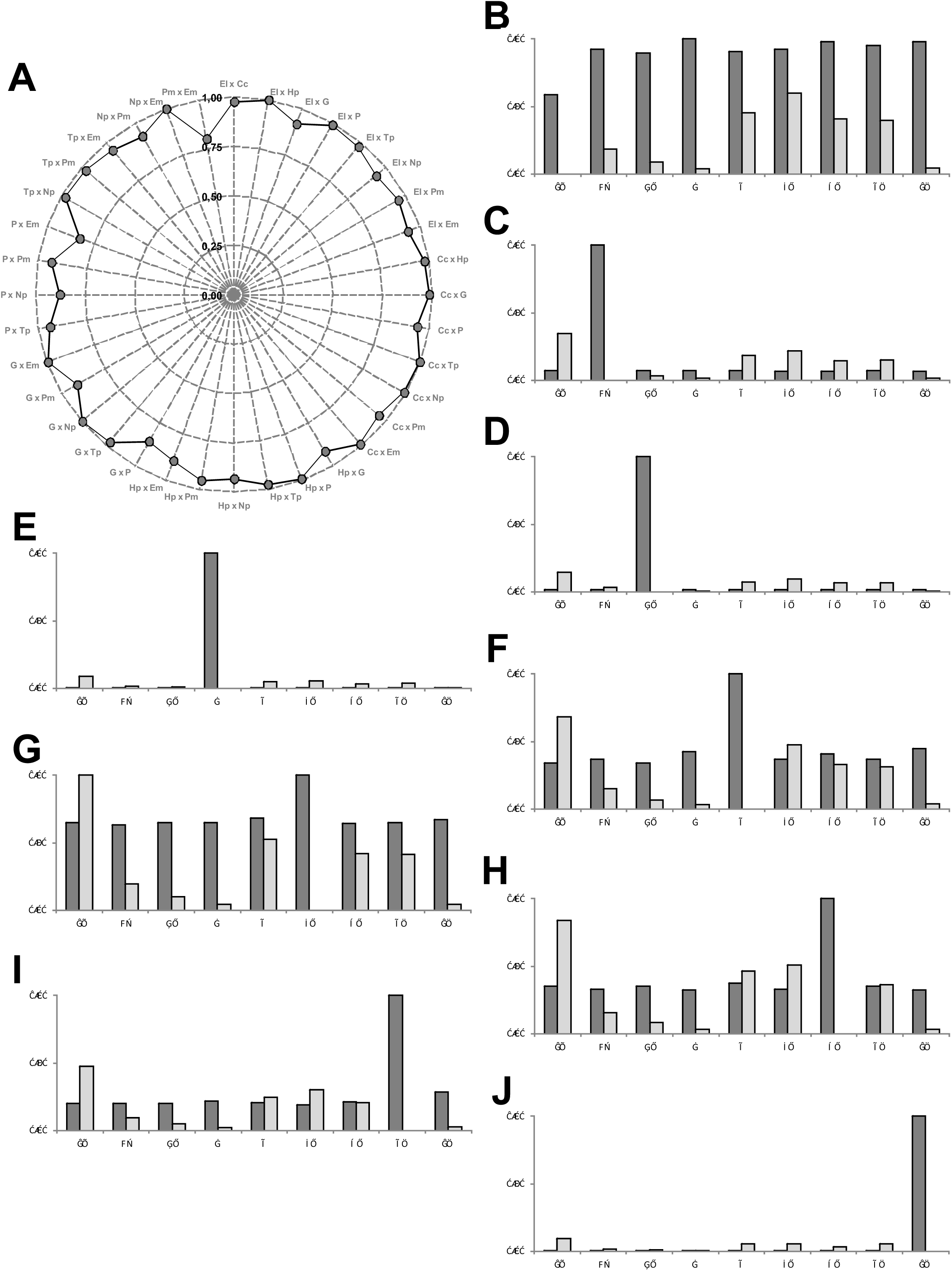
(A) Isotopic niche overlap (*O*). Intraspecific competition (*IC*) and Interespecific competition (*SC*) of extant megamammals from Africa, (B) *L. africana*, (C) *E. quagga*, (D) *D. bicornis*, (E) *C. simum*, (F) *C. taurinus*, (G) *S. caffer*, (H) *K. ellipsiprymnus*, (I) *O. beisa*, (J) *G. camelopardalis*, (K) *H. amphibius*.

Niche overlap between grazers and mixed-feeders (*L. africana* and *D. bicornis*) was moderate (*O* = 0.49-0.53), mainly because these taxa fed more in fruits (*p*_*i*_ = 0.26-0.34). However, although they fed little on C_4_ grass (*p*_*i*_ = 0.21-0.24), if these species try to compete for grasses, they were much more competitive than grazers (Table 5; Figure 6; *e.g.* Boer *et al*., 2015). *Hippopotamus amphibius* had a great consumption of C_4_ grass (*p*_*i*_ = 0.58), and presents a moderate niche overlap with *C. simum* and *C. taurinus* (*O* = 0.54) and a high niche overlap with remain grazers (*O* = 0.84-0.87), being a little better competitor for this kind of resource (Table 5; Figure 6).

*Giraffa camelopardalis* was the only browser evaluated, presenting a moderate niche overlap with *L. africana* and *D. bicornis* (*O* = 0.40 and 0.63, respectively), being a weak competitor with these taxa (Table 5), and had a low niche overlap with grazers (*O* = 0.01-0.33).

The best competitor species in Africa mammal fauna through interspecific competition index (Table 5; Figure 6) was *Loxodonta africana*, followed by *D. bicornis*, as it was already observed in experimentation (Landman *et al.*, 2013).

Based in these observations on Africa meso-megaherbivores fauna, we can make some assumptions for Pleistocene meso-megaherbivores fauna from Poço Redondo, Sergipe State. First of all, we note that niche overlap was high for all taxa (*O* = 0.80-1.00), mainly because mixed feeders taxa had a great consumption of C_4_ grass (*p*_*i*_ = 0.59-0.80).

*N. platensis* (*w* = 6,265 kg) was the unique grazer, and as in grazers from Africa, the intraspecific competition was higher, and apparently more important, than interspecific competition (Table 5; Figure 7).

**Figure 7.**
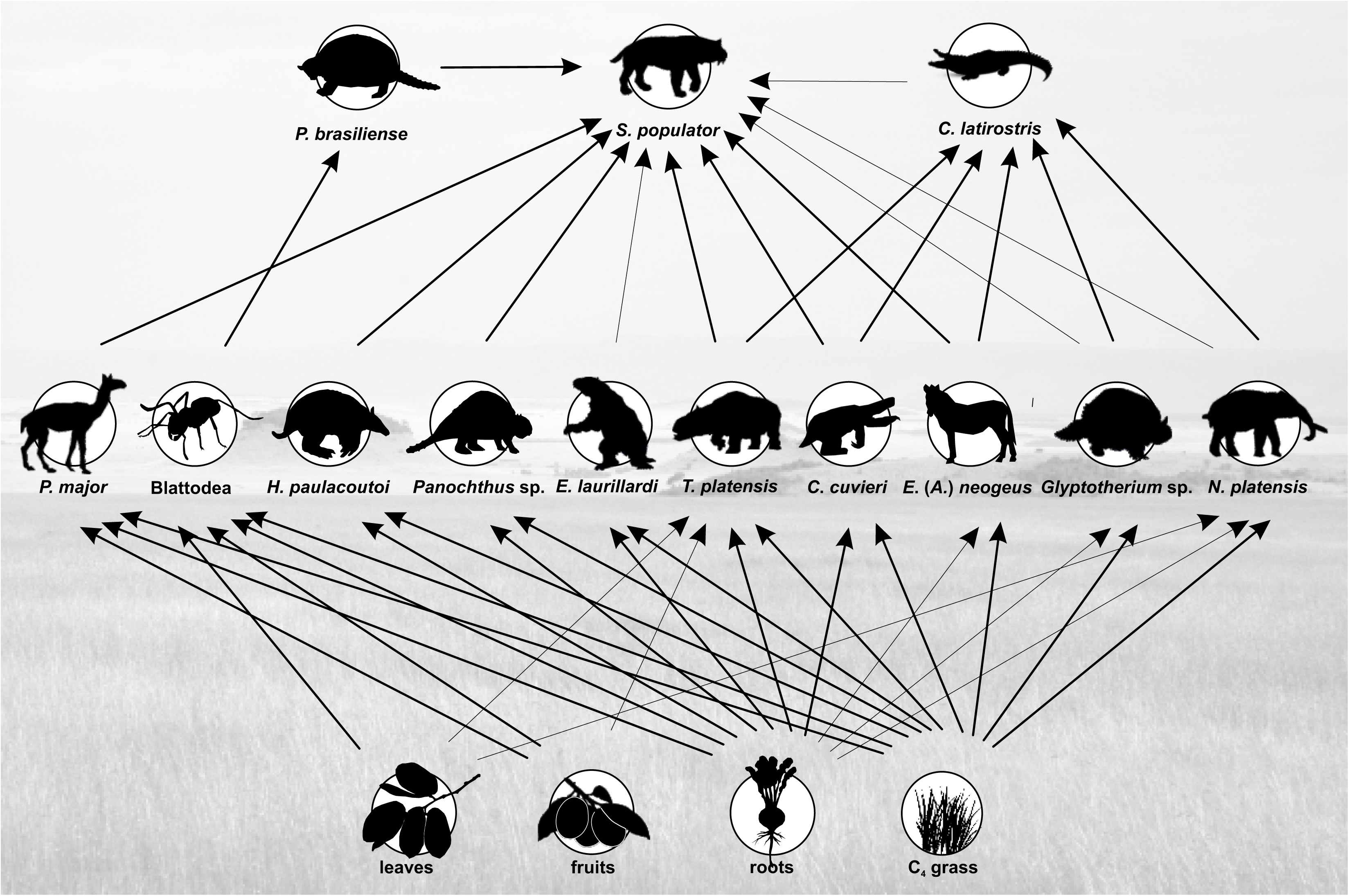
(A) Isotopic niche overlap (*O*). Intraspecific competition (*IC*) and Interespecific competition (*SC*) of pleistocenic megamammals from Sergipe, Brazil, (B) *E. laurillardi*, (C) *C. cuvieri*, (D) *H. paulacoutoi*, (E) *Glyptotherium* sp., (F) *Panochthus* sp., (G) *T. platensis*, (H) *N. platensis*, (I) *P. major*, (J) *E. (A.) neogeus*.

The mixed-feeder guild was composed of eight taxa, two megaherbivores (*E. laurillardi*, *w* = 3,416 kg; *T. platensis*, *w* = 1,770 kg) and six mesoherbivores (*H. paulacoutoi*, *w* = 120 kg; *P. major*, *w* = 285 kg; *C. cuvieri*, *w* = 777 kg; *E.* (*A.*) *neogeus*, *w* = 420 kg; *Glyptotherium* sp., *w* = 710 kg; *Panochthus* sp., *w* = 785 kg).

As in Africa all mixed-feeders were better competitors than grazers, presenting better *IC* (Table 5). The better competitors in the Late Pleistocene of Poço Redondo, *E. laurillardi* (Table 5, Figure 7), followed by *Toxodon platensis*.

### 3.5. Paleoenvironmental reconstruction

The available isotopic data from extant megamammals from Kenya and Tanzania (Table 3; Figure 3A) provide us a good portrait of meso-megaherbivores fauna from Africa, with a predominance of grazer species with high consumption of C_4_ plants (*p*_*i*_ = 92-100 %; Table 3), and where even mixed-feeder and browser species had a high consumption of grass (*p*_*i*_ = 21-58 %). Roots consumption was low (*p*_*i*_ = 1-24 %; values for *H. amphibius* represents C_3_ aquatic plants). Based on *Loxodonta africana* δ^18^O_water_ we estimate a Mean Annual Temperature - MAT around 29±3 °C for Africa.

Using regressions proposed by Coe *et al.* (1976) for our Africa mammal assemblage, we found similar values of Energy Expenditure, Biomass, Production and Annual Precipitation for assemblies in wildlife areas in Kenya and Tanzania (Table 6), which show us that we have enough information to estimate the same measures in Late Pleistocene meso-megaherbivores fauna from Poço Redondo, Sergipe, Brazil.

**Table 6.** Biomass, expendidure energy (EE), secondary production and annual precipitation in localities from Africa in comparison to our meso-megamammal assembly from Africa and Pleistocene of Sergipe, Brazil. **References:** ^(1)^ Coe *et al.* (1976); ^(2)^ Our data.

| Locality | Biomass (Kg/km<sup>2</sup>) | EE (Kj/Km<sup>2</sup>.h) | Sec. Production (Kg/Km<sup>2</sup>.yr) | AP (mm) |
| --- | --- | --- | --- | --- |
| Amboseli, Kenya<sup>(1)</sup> | 4,848 | 22,970 | 934 | 350 |
| Nairobi National Park, Kenya<sup>(1)</sup> | 4,824 | 24,728 | 1,008 | 844 |
| Loliond Controld Area, Tanzania<sup>(1)</sup> | 5,423 | 26,962 | 1,134 | 784 |
| Serengeti National Park, Tanzania<sup>(1)</sup> | 8,352 | 43,063 | 1,743 | 803 |
| Ruaha National Park, Tanzania<sup>(1)</sup> | 3,909 | 12,474 | 364 | 625 |
| Meso-megamammals from Africa<sup>(2)</sup> | 4,485 | 29,666 | 1,278 | 565-845 |
| Meso-megamammals from, Sergipe, Brazil<sup>(2)</sup> | 4,842 | 37,410 | 1,250 | 585-832 |

In the Late Pleistocene of Sergipe the available data of meso-megaherbivores fauna shows a mammal assemblage composed of mixed-feeders and grazer species (Table 3). They fed more on C_4_ grass (*p*_*i*_ = 59-94 %) than the extant fauna from Africa, which could indicate that they lived in a more open and drier environments. Another evidence is that this fauna fed more on roots than Africa fauna (*p*_*i*_ = 6-41 %; Table 3), which in combination to the high values of δ^18^O_water_ found in *N. platensis*, suggests a drier environment than in Africa, with high MAT (37±6 °C, Table 6).

Biomass (4,733 kg/km^2^) and Secondary Production (1,192 kg/km^2^.yr) were similar to that found in wildlife areas from Kenya and Tazania (Table 6), however Energy Expenditure was high in Poço Redondo (35,049 kj/km^2^.h). Another difference, although Poço Redondo was hotter than Africa nowadays, is that it had probably a similar Annual Precipitation, varying to our estimations between 585 to 832 mm. If the meso-megaherbivores lived in a drier environment, this could explain why they fed more on roots, as this is one of the main components of Net Primary Production (NPP) in Seasonal Dry Forests (Jaramillo *et al.*, 2011 and references therein).

## Conclusion

In this paper we investigated isotopic data for Africa meso-megamammals to help undestanding better the paleoecology of meso-megamammals from Late Pleistocene of Poço Redondo (Sergipe, Brazil). First of all, we estimated the weights of these taxa, and noticed that in Poço Redondo the vertebrate assemblage was composed at least by one megacarnivore (*S. populator*, *w* = 315 kg), one small carnivore (*C. latirostris*, *w* = ~60 kg), one omnivore (*P. brasiliense*, *w* = 38 kg), six mesoherbivores (*H. paulacoutoi w* = 120 Kg; *P. major*, *w* = 285 kg; *E.* (*A.*) *neogeus*, *w* = 420 kg; *Glyptotherium* sp., *w* = 710 kg; *C. cuvieri*, *w* = 777 Kg; and *Panochthus* sp., *w* = 785 kg;) and three megaherbivores (*T. platensis*, *w* = 1,770 kg; *E. laurillardi*, *w* = 3,416 kg; and *N. platensis*, *w* = 6,265 kg).

The herbivore fauna had a high consumption of C_4_ grass, belonging to two guilds: grazers (*p*_*i*_C_4_ grass > 90%; *N. platensis*) and Mixed Feeders (*Glyptotherium* sp; *E. neogeus*; *T. platensis*; *H. paulacoutoi*; *Panochthus* sp.; *C. cuvieri*; *E. laurillardi*; and *P. major*).

*Smilodon populator* could fed on meso-megaherbivores weighting between 285 kg to 6,300 kg (*p*_*i*_ = 67 %), eventually could have hunted *C. latirostris* (*p*_*i*_ = 10 %), and we suggests that acted as scavengers feeding in carcass of *Pachyarmatherium brasiliense* (*p*_*i*_ = 11 %) and *H. paulacoutoi* (*p*_*i*_ = 12 %). Beside this, we suggest that *C. latirostris* could act as a scavenger, feeding on dead corpses of these meso-megamammals, among which *N. platensis* (*p*_*i*_ = 44 %) was a great contributor based on its isotopic signature.

Through niche overlap (*O*) and intra-interspecific (*IC*/*SC*) indexes we noticed that in meso-megamammals of Africa that mixed-feeders are better competitors than grazers, allowing to suggest that *Eremotherium laurillardi* (*w* = 3,416 kg; *B*_*A*_ = 0.23) and *Toxodon platensis* (*w* = 1,770 kg; *B*_*A*_ = 0.24), respectively, were the best resources competitors in the Late Pleistocene mammal assemblage of Poço Redondo, indicating that large weights are important to determine a good competitor.

Finally, we suggest that the meso-megamammals from the Late Pleistocene of Poço Redondo, Sergipe, lived in an open and dry environment similar to that found nowadays in Africa, with similar biomass and annual precipitation, but hotter and with a higher energy expenditure for the megafauna.

## Acknowledgements

To CNPq by financial support through Universal project (process 404684/2016-5); To Dr. Alexandre Liparini (Laboratório de Paleontologia/UFS) and Dr. Castor Cartelle (PUC/MG) for allow measurements in fossils collection that they are responsible for; To biologists Verônica Gomes and Lais Alves for help in Solver (Excel) analysis; And for Dr. Hermínio Araujo Jr. and M.Sc. Lidiane Asevedo for critical evaluation of manuscript.

---

**Supplementary Table 1.**
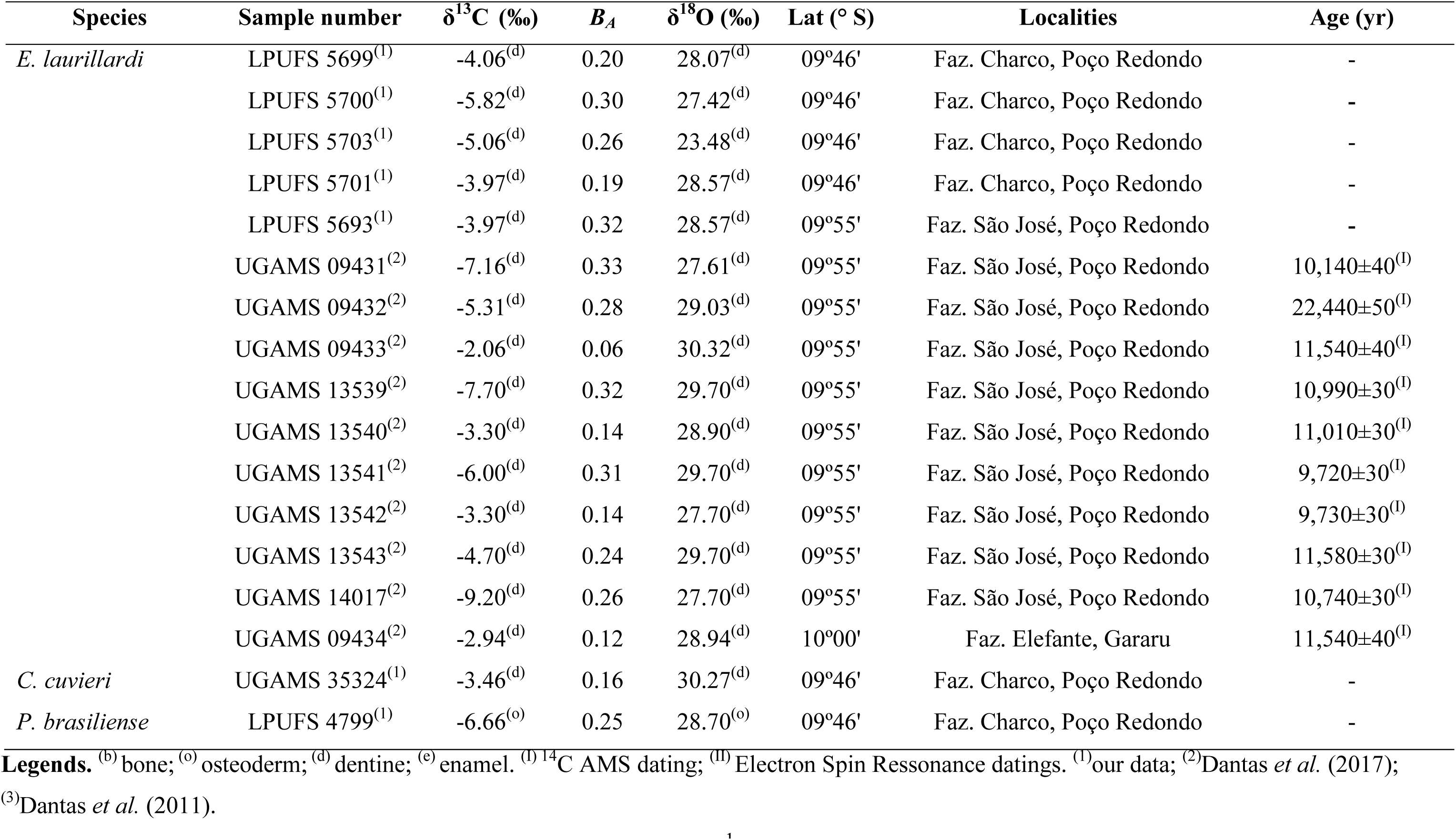
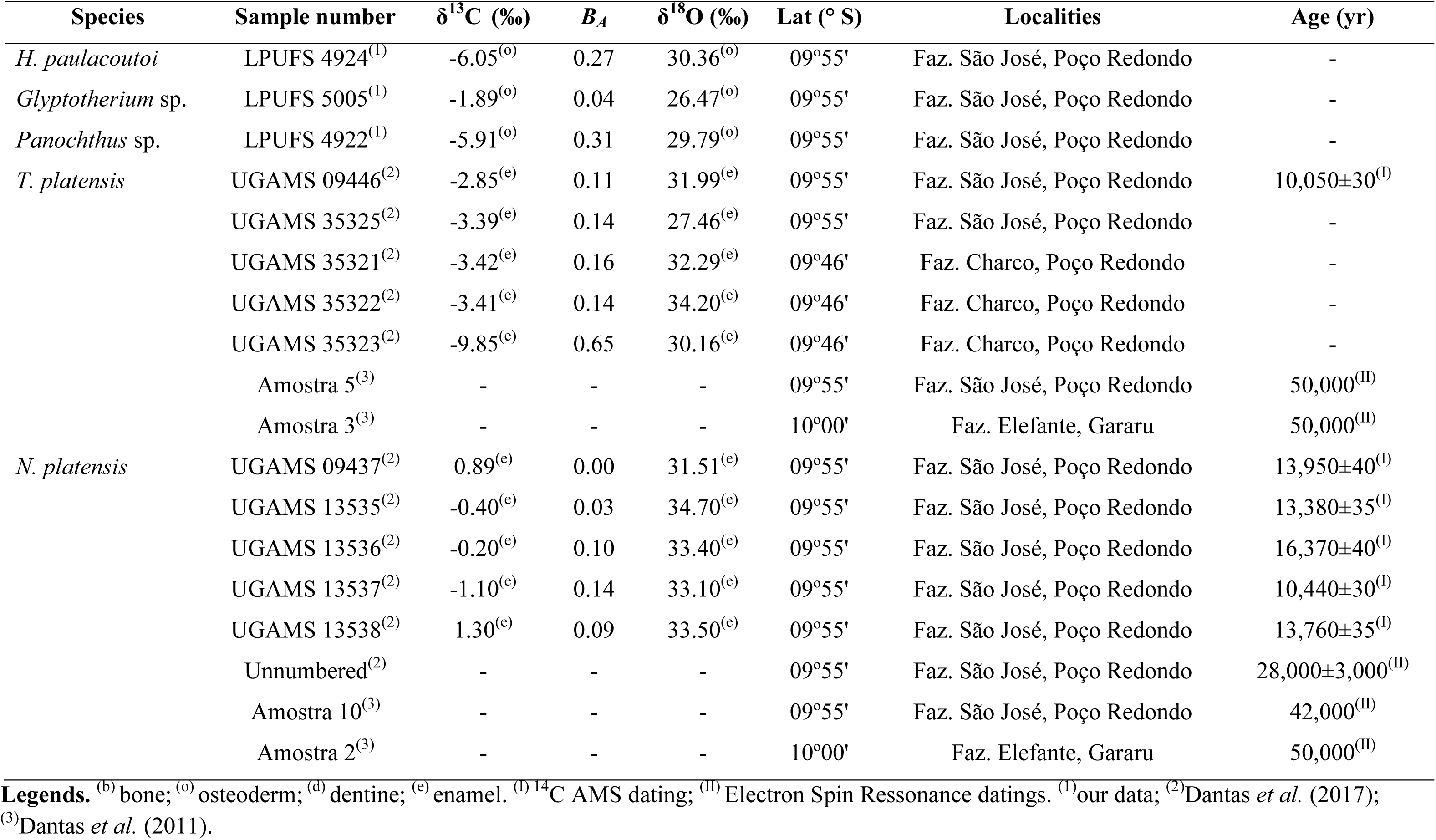
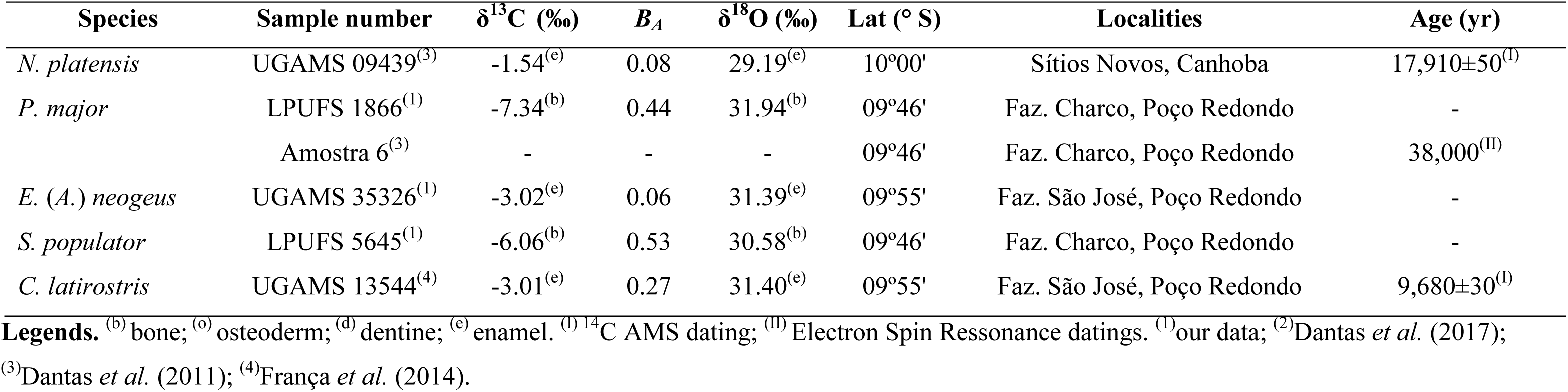
Carbon (in VPDB) and oxygen (in VSMOW) isotopic values and available datings for ten extinct late Pleistocene vertebrate taxa from Sergipe, Brazil.

**Supplementary Table 2.** Weight estimation for several taxa of Pleistocenic megafauna from Brazilian Intertropical Region.

| Taxa | humerus |  | femur |  | Weight (Kg) | Localities |
| --- | --- | --- | --- | --- | --- | --- |
|  | Sample | Circ. | Sample | Circ. |  |  |
| *E. laurillardi* | LPUFS 2101 | 350.00 |  |  |  | Monte Alegre/SE<sup>(1)</sup> |
|  |  |  | LPUFS | 740.00 |  | Monte Alegre/SE<sup>(1)</sup> |
|  |  |  | LPUFS 2264 | 658.00 |  | Poço Redondo/SE<sup>(1)</sup> |
| mean |  | 350.00 |  | 699.00 | 3,416.18 |  |
| *C. cuvieri* | MCL 22470/02 | 211.00 |  |  |  | Nova Redenção/BA<sup>(1)</sup> |
|  | MCL 22473 | 240.00 |  |  |  | Nova Redenção/BA<sup>(1)</sup> |
|  | MCL 22475 | 216.00 |  |  |  | Nova Redenção/BA<sup>(1)</sup> |
|  |  |  | MCL 22500 | 325.00 |  | Nova Redenção/BA<sup>(1)</sup> |
|  |  |  | MCL 22394/08 | 394.00 |  | Nova Redenção/BA<sup>(1)</sup> |
| mean |  | 222.33 |  | 359.50 | 777.73 |  |
| *P. brasiliense* | MCC 996-V | 59.66 |  |  |  | Baraúna/RN<sup>(2)</sup> |
|  |  |  | MCC 1133-V | 153.86 |  | Baraúna/RN<sup>(2)</sup> |
| mean |  | 59.66 |  | 153.86 | 38.03 |  |
| *Panochthus* sp. | UESB 318PV172 | 136.00 |  |  |  | Anagé/BA<sup>(1)</sup> |
|  | MN 2964-V | 120.26 |  |  |  | Taperoá/PB<sup>(3)</sup> |
|  |  |  | MN 2760-2V | 239.08 |  | Taperoá/PB<sup>(3)</sup> |
| mean |  | 128.13 |  | 239.08 | 783.99 |  |

**Supplementary Table 2 (continuation).**
| Taxa | humerus |  | femur |  | Weight (Kg) | Localities |
| --- | --- | --- | --- | --- | --- | --- |
|  | Sample | Circ. | Sample | Circ. |  |  |
| *Glyptotherium* sp. | MCC 1087V | 102.05 |  |  |  | Baraúna/RN<sup>(2)</sup> |
|  |  |  | MCC 1560V | 252.30 |  | Baraúna/RN<sup>(2)</sup> |
| mean |  | 102.05 |  | 252.30 | 711.28 |  |
| *H. paulacoutoi* | MCL 501/02 | 123.00 |  |  |  | Jacobina/BA<sup>(1)</sup> |
|  |  |  | MCL 501/08 | 155.00 |  | Jacobina/BA<sup>(1)</sup> |
| mean |  | 123.00 |  | 155.00 | 120.64 |  |
| *N. platensis* | MHNT-VT 2035 | 523.16 |  |  |  | São Bento do Una/PE<sup>(4)</sup> |
|  | MHNT-VT 2036 | 386.95 |  |  |  | São Bento do Una/PE<sup>(4)</sup> |
|  | MHNT-VT 2037 | 338.78 |  |  |  | São Bento do Una/PE<sup>(4)</sup> |
|  |  |  | MHNT-VT 1138 | 510.44 |  | São Bento do Una/PE<sup>(4)</sup> |
|  |  |  | MHNT-VT 2031 | 325.08 |  | São Bento do Una/PE<sup>(4)</sup> |
|  |  |  | MHNT-VT 2032 | 274.40 |  | São Bento do Una/PE<sup>(4)</sup> |
| mean |  | 416.30 |  | 369.97 | 6,266.18 |  |
| *T. platensis* | LPUFS 2188 | 265.00 |  |  |  | Poço Redondo/SE<sup>(1)</sup> |
|  |  |  | LPUFS 5691 | 230.00 |  | Poço Redondo/SE<sup>(1)</sup> |
| mean |  | 265.00 |  | 230.00 | 1,771.60 |  |

**Supplementary Table 2 (continuation).**
| Taxa | humerus |  | femur |  | Weight (Kg) | Localities |
| --- | --- | --- | --- | --- | --- | --- |
|  | Sample | Circ. | Sample | Circ. |  |  |
| *P. major* | UESB 318PV172 | 136.00 |  |  |  | Anagé/BA<sup>(1)</sup> |
|  |  |  | UESB 318PV173 | 117.00 |  | Anagé/BA<sup>(1)</sup> |
| Mean |  | 136.00 |  | 117.00 | 283.54 |  |
| *E. (A.) neogeus* | MCL 6212 | 140.00 |  |  |  | Ourolândia/BA<sup>(1)</sup> |
|  |  |  | MCL 6229 | 152.00 |  | Ourolândia/BA<sup>(1)</sup> |
| mean |  | 140.00 |  | 152.00 | 419.36 |  |
| *S. populator* | MCL 7187/48 | 137.00 |  |  |  | Campo Formoso/BA<sup>(1)</sup> |
|  | MCL 2998 | 151.00 |  |  |  | Ourolândia/BA<sup>(1)</sup> |
|  |  |  | MCL 7160 | 124.00 |  | Jacobina/BA<sup>(1)</sup> |
|  |  |  | MCL 7161 | 114.00 |  | Jacobina/BA<sup>(1)</sup> |
| mean |  | 144.00 |  | 119.00 | 315.19 |  |
| *M. tridactyla* | MCL 1602/06 | 72.00 |  |  |  | Morro do Chapéu/BA<sup>(1)</sup> |
|  | MCN-M 99 | 100.00 |  |  |  | Morro do Chapéu/BA<sup>(1)</sup> |
|  |  |  | MCL 1602/07 | 83.00 |  | Belo Horizonte (Zoo)<sup>(1)</sup> |
|  |  |  | MCN-M 99 | 74.00 |  | Belo Horizonte (Zoo)<sup>(1)</sup> |
| mean |  | 86.00 |  | 78.50 | 34.86 |  |

**Supplementary Table 2 (continuation).**
|  |  |  |  |  |  |  |
| --- | --- | --- | --- | --- | --- | --- |
| *P. maximus* | MACN s/n | 98.91 |  |  |  | locality unavailable<sup>(5)</sup> |
|  |  |  | MACN s/n | 70.02 |  | locality unavailable<sup>(5)</sup> |
| mean |  | 98.91 |  | 70.02 | 43.12 |  |
| *T. tetradactyla* | LEG 0644 | 39.00 |  |  |  | Campo Formoso/BA<sup>(1)</sup> |
|  | LEG s/n | 44.00 |  |  |  | Ituaçu/BA<sup>(1)</sup> |
|  |  |  | LEG 0656 | 31.00 |  | Campo Formoso/BA<sup>(1)</sup> |
|  |  |  | LEG s/n | 39.00 |  | Ituaçu/BA<sup>(1)</sup> |
| mean |  | 41.50 |  | 35.00 | 4.57 |  |

**References.** <sup>(1)</sup> our data; <sup>(5)</sup> Fariña & Vizcaino (1997).

**Supplementary Table 3.**
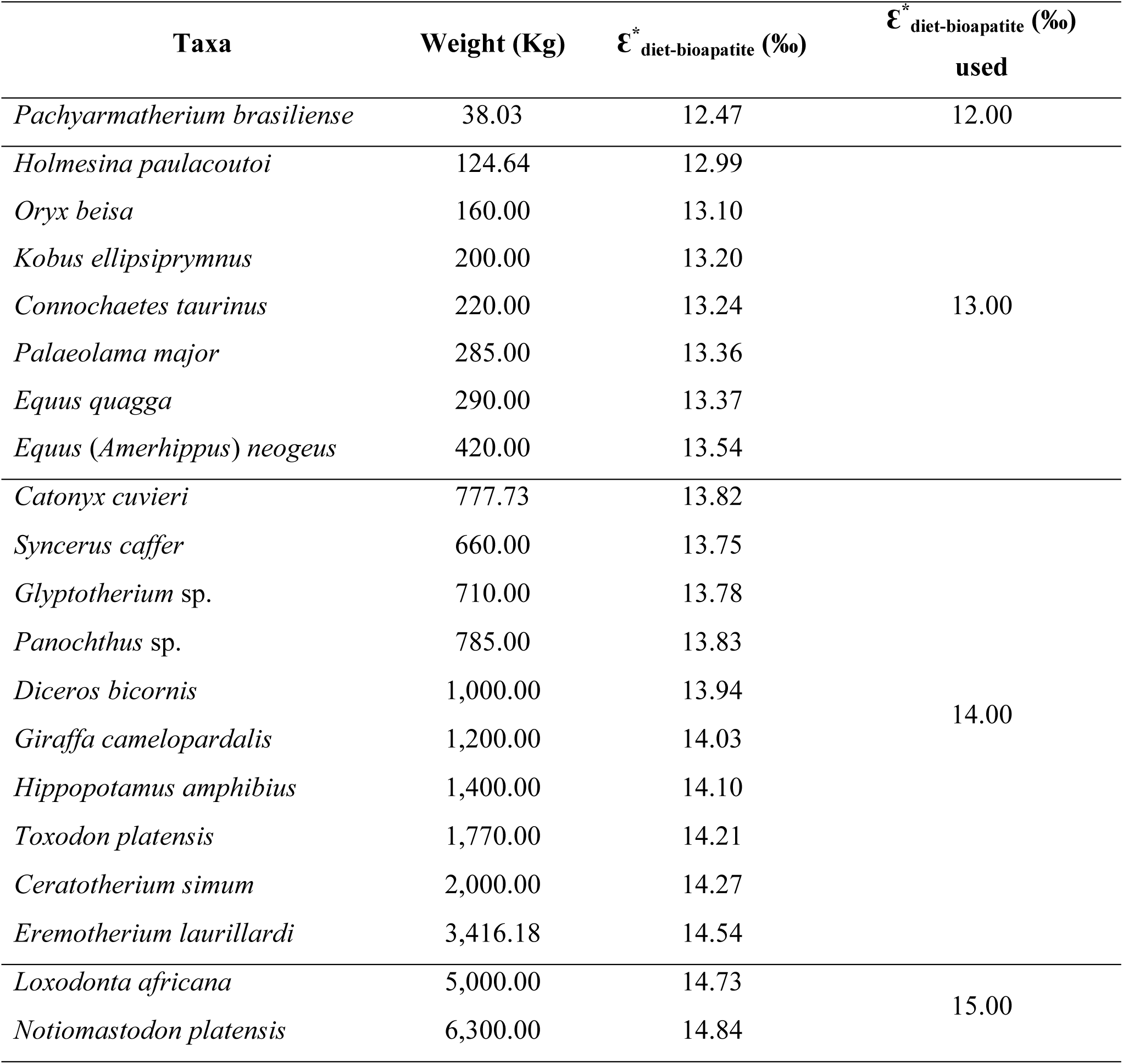
Estimated carbon (δ^13^C) diet-bioapatite enrichment (ε^*^ _diet-bioapatite_) in herbivores from Africa and Sergipe.

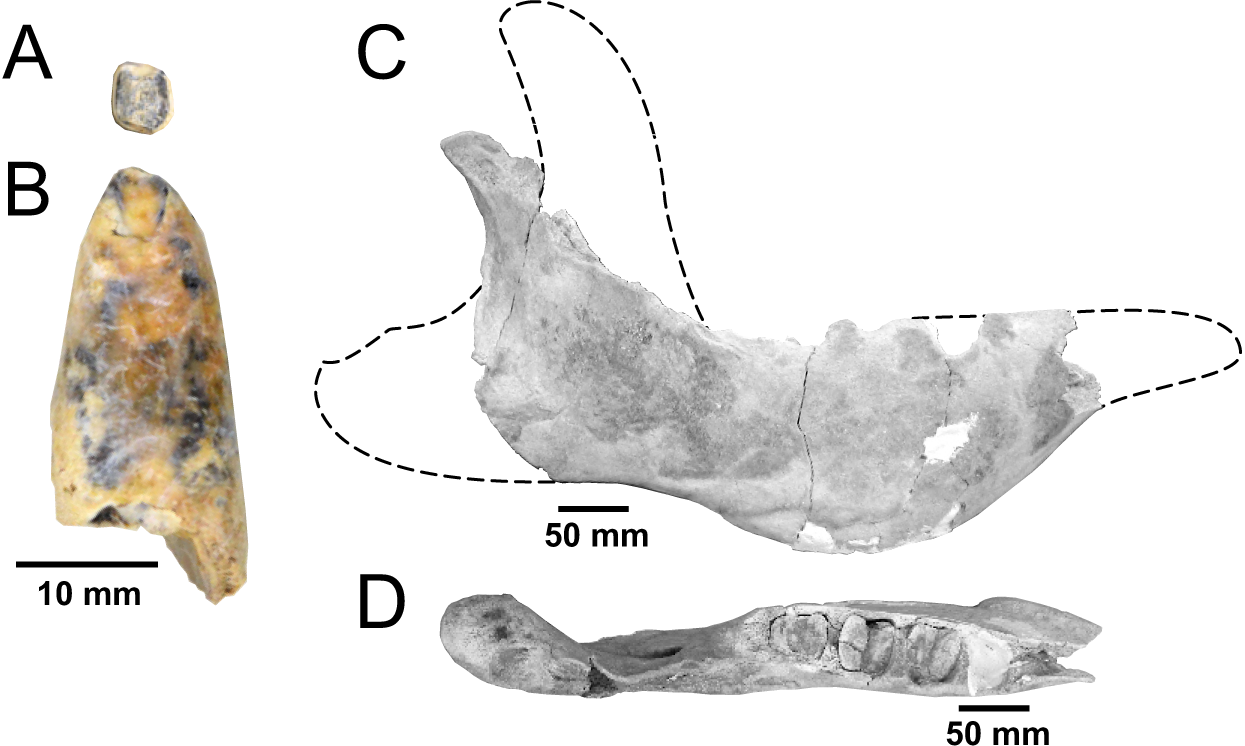

